# No evidence for an effect of M1 cTBS on schema-mediated motor sequence learning

**DOI:** 10.64898/2026.03.12.711304

**Authors:** S. Reverberi, K. Cuypers, B.R. King, G. Albouy

**Affiliations:** Department of Movement Sciences, Movement Control and Neuroplasticity Research Group, KU Leuven, Leuven, Belgium; Defitech Chair of Clinical Neuroengineering, INX and BMI, EPFL Valais, Clinique Romande de Réadaptation, Sion, Switzerland; REVAL—Rehabilitation Research Center, Faculty of Rehabilitation Sciences, University of Hasselt, Diepenbeek, Belgium; Department of Health and Kinesiology, College of Health, University of Utah, Salt Lake City, UT, USA

**Keywords:** motor learning, schema, TMS, M1, memory consolidation

## Abstract

The availability of a pre-existing cognitive-motor schema accelerates the learning of novel motor information. The encoding of a novel schema-compatible, compared to-incompatible, motor sequence was recently shown to be supported by the left primary motor cortex (M1). However, causal evidence for the role of M1 in schema-mediated motor learning is currently lacking. In the current study, we aimed to address this knowledge gap by transiently disrupting M1 using inhibitory continuous theta burst stimulation (cTBS). Forty-eight young healthy participants learned a bimanual motor sequence task (cognitive-motor schema). Twenty-four hours later, they learned a novel sequence whose ordinal schematic structure was compatible with that learned on the previous day. To provide causal evidence for a role of M1 on such schema-mediated motor learning, we applied either cTBS or sham stimulation to the left M1 immediately prior to encoding the schema-compatible novel sequence. Electromyography results showed no evidence for an effect of left M1 cTBS on corticospinal excitability as measured with motor-evoked potentials. Similarly, behavioral results indicated no significant effect of cTBS on subsequent schema-mediated motor sequence learning. Altogether, the present data do not provide evidence for a causal role of the left M1 in schema-mediated motor sequence learning.

## 1. Introduction

Memory consolidation, i.e., the stabilization of initially labile memory traces, is traditionally described as a long process occurring over a timescale ranging from days to years (Dudai et al., 2015). Recent research, however, suggests that memory consolidation can be accelerated by the availability of a pre-existing schema. Schemas are defined as structures of associative knowledge, formed through the repeated encounter of related items or experiences, and consisting of their abstracted, conceptual representation (Lewis & Durrant, 2011). When novel memories to be encoded are compatible with a pre-established schema, they are rapidly consolidated into long-term memory (Lewis & Durrant, 2011). This schema-mediated acceleration of memory consolidation was first demonstrated in rats through the rapid learning of novel flavor-location associations consistent with a previously learned flavor-location schema (Tse et al., 2007, 2011), and then in humans through the rapid learning of novel, schema-compatible associative knowledge (e.g., object-scene, item-color, noun-adjective associations; Bein et al., 2015; Cycowicz et al., 2008; van Kesteren et al., 2013). Perceptual learning has also been described to benefit from previously acquired schema, as earlier research suggests that new melodies that conform to a known tonal schema are more easily learned than non-conformant memories (Durrant et al., 2015). Recently, the schema model of memory consolidation was extended to the motor memory domain, as the learning of a novel sequence of finger movements was demonstrated to be significantly enhanced if its ordinal schema (i.e., the temporal position of the movements in the sequence stream) was compatible with that of a previously learned motor sequence (King et al., 2019).

At the brain level, declarative memory schemas are described to be stored in the neocortex, with the medial prefrontal cortex (mPFC) being involved in the fast encoding of schema-compatible information (Lewis & Durrant, 2011; van Kesteren et al., 2012, 2013). Novel material that is compatible with an acquired schema is thought to bypass short-term hippocampal storage and thus be directly assimilated into the neocortical schema (Lewis & Durrant, 2011; Tse et al., 2007; van Kesteren et al., 2012). Causal evidence supporting such view comes from previous transcranial magnetic stimulation (TMS) and lesion studies. Specifically, TMS has been used to investigate the role of the mPFC in schema-related processes using the Deese–Roediger–McDermott (DRM) task, during which participants are taught lists of words that belong to the same semantic neighborhood (e.g., “tired”, “night”, “pillow”). This procedure results in the development of a schema that can then induce the false recall of critical lures, i.e. unstudied schema-congruent words (e.g. “sleep”, Roediger & McDermott, 1995). Disruptive continuous theta-burst stimulation (cTBS) delivered to the mPFC, but not to the vertex, prior to task performance was shown to significantly reduce false recall of critical lures (Berkers et al., 2017). Similarly, cTBS of the mPFC prior to the encoding of an emotional DRM task significantly reduced false recognition of negative lures, compared to control 5Hz repetitive TMS (Bovy et al., 2020). Additionally, lesions to the ventromedial PFC were shown to result in impaired classification of stimulus congruency with schema (Ghosh et al., 2014) and reduced false recall of critical lures (Warren et al., 2014). Finally, vmPFC lesions were also shown to result in a lack of schema benefit for recognition of congruent items (Spalding et al., 2015). Overall, these studies provide causal evidence for the involvement of the mPFC in schema-mediated declarative memory encoding.

In contrast to the declarative memory domain, the neural processes supporting schema-facilitated non-declarative learning have only been examined more recently. Evidence from the motor memory domain suggests that the left primary motor cortex (M1) is recruited during the encoding of a novel schema-compatible (as compared to incompatible) motor sequence (Reverberi et al., 2025). Multivoxel activity patterns in M1 were also altered around novel movements when they were learned in a context that was compatible – as compared to incompatible – with the previously acquired cognitive motor-schema (Reverberi et al., 2025). The results of this study therefore suggest a role for the left M1 in the rapid integration of novel information into a compatible motor schema. However, causal evidence for this role is lacking. In the current study, that was pre-registered with Open Science Framework (https://osf.io/qe5vc), we therefore aimed to provide causal evidence for the role of the left M1 in schema-mediated integration of novel information into motor memory. To this end, we applied cTBS to the left M1 prior to the learning of a novel schema-compatible motor sequence. Briefly, participants first learned a bimanual motor sequence task, from which a schematic ordinal structure representing the binding of each movement to its temporal position in the sequence is believed to be extracted (ordinal-based cognitive-motor schema, Dolfen et al., 2024; King et al., 2019; Reverberi et al., 2025). Approximately 24 hours later, participants received either cTBS or sham stimulation of the left M1 immediately before learning a novel, schema-compatible sequence that contained both *learned* and *novel* movement transitions (i.e., transitions already present in the sequence learned the previous day, and transitions introduced on the second day, respectively). This design allowed us to examine the effect of inhibitory stimulation of M1 on both the retention of previously learned movements and the learning of new movements that are performed in a context that is compatible with previous schema. Based on previous work showing that disruptive TMS delivered to M1 24 hours after initial learning did not affect finger-tapping performance at a subsequent retest (Hotermans et al., 2008), we hypothesized that cTBS – as compared to sham stimulation – over the left M1 would not affect performance on learned movement transitions. In contrast, based on (1) previous TMS work showing that cTBS administered to M1 before learning a new motor task impaired subsequent sequence learning and 30-min recall performance (Rosenthal et al., 2009; Wilkinson et al., 2010, 2015), and (2) our previous neuroimaging work suggesting a critical role for the left M1 in schema-mediated integration of new motor information into memory (Reverberi et al., 2025), we expected cTBS of M1 to impair performance on the novel transitions, reflecting a disruption of the integration of new movements into compatible schema.

## 2. Methods

Data collection and analysis plans for this study were pre-registered with the Open Science Framework (https://osf.io/qe5vc). Data collection and primary statistical analyses did not deviate from the original pre-registered plan. Any additional (non-pre-registered) analyses are labelled in the manuscript as exploratory.

### 2.1 Participants

Fifty-one healthy young volunteers (mean age: 24, range: 19-30 years old) were recruited from Leuven and the surrounding area. The inclusion criteria for participation were: 1) right-handed, as assessed with the Edinburgh Handedness Inventory (Oldfield, 1971), 2) no prior extensive training with a musical instrument requiring dexterous finger movements (e.g., piano, guitar) or as a professional typist, 3) free of medical, neurological, psychological, or psychiatric conditions, including depression and anxiety as assessed by the Beck’s Depression and Anxiety Inventories (Beck et al., 1988, 1996), 4) no indications of abnormal sleep, as assessed by the Pittsburgh Sleep Quality Index (PSQI (Buysse et al., 1989)); 5) not considered extreme morning or evening types, as quantified with the Horne & Ostberg chronotype questionnaire (Horne & Ostberg, 1976), 6) free of psychoactive and sleep-influencing medications, 7) non-smokers, 8) not having completed trans-meridian trips or worked night shifts in the month prior to participation, and 9) presenting no contra-indications to TMS procedures. Participants gave written informed consent prior to the start of the study that was approved by the local ethics committee, and they received a monetary compensation for their participation. Participants were pseudo-randomly assigned to the active stimulation (STIM) or the sham stimulation (SHAM) group to achieve an approximately equal gender ratio between groups (see Table 1).

**Table 1:** Participant characteristics, sleep and vigilance scores for each experimental group. Group means are presented, with standard deviation in parentheses. Average sleep duration for the 3 nights prior to Session 1 (S1) was assessed via self-reported sleep diaries. Sleep duration for the night prior to Session 2 (S2) was assessed via visual inspection of actigraphy data extracted from wrist-mounted device. **A.** Statistical differences between groups were assessed with independent samples t-tests. Degrees of freedom = 1,45 for Average sleep duration, 3 nights prior to S1 and Sleep duration, night prior S2; degrees of freedom = 1,46 for all other variables. **B,C**. Statistical differences between groups were assessed with 2 (session) x 2 (group) ANOVAs (degrees of freedom = 1,46 for all). S x G = session x group. B: BAI = Beck Anxiety Inventory (Beck et al., 1988); BDI = Beck Depression Inventory (Beck et al., 1996); PSQI = Pittsburgh Sleep Quality Index (Buysse et al., 1989); SMS = St. Mary’s Hospital Sleep Questionnaire (Ellis et al., 1981); PVT = Psychomotor Vigilance Task (Dinges & Powell, 1985); SSS = Stanford Sleepiness Scale (Hoddes, 1972).

| A. Variable | Group mean (SD) |  | t | p | Cohen's d |  |
| --- | --- | --- | --- | --- | --- | --- |
|  | STIM | SHAM |  |  |  |  |
| N | 24 | 24 |  |  |  |  |
| Female (n) | 15 | 16 |  |  |  |  |
| Age (years) | 24.0 (3.0) | 24.2 (2.9) | -0.15 | 0.88 | -0.042 |  |
| BAI score | 2.1 (2.2) | 3.6 (3.9) | -1.69 | 0.10 | -0.487 |  |
| BDI score | 4.7 (5.7) | 4.4 (5.3) | 0.21 | 0.26 | 0.061 |  |
| Handedness score | 87.7 (12.9) | 83.8 (13.4) | 1.05 | 0.30 | 0.302 |  |
| PSQI score | 3.3 (1.9) | 3.0 (1.7) | 0.65 | 0.52 | 0.186 |  |
| Chronotype score | 51.5 (7.5) | 50.0 (8.7) | 0.64 | 0.53 | 0.185 |  |
| Average sleep duration, 3 nights prior to S1 (hours) | 8.2 (0.6) | 8.5 (0.7) | -1.86 | 0.07 | -0.541 |  |
| Sleep duration, night prior to S2 (hours) | 8.3 (1.2) | 8.4 (1.5) | -0.24 | 0.81 | -0.069 |  |
| S1 SMS duration (hours) | 7.9 (0.8) | 8.1 (0.8) | -0.82 | 0.42 | -0.236 |  |
| S2 SMS duration (hours) | 8.1 (1.1) | 8.2 (1.2) | -0.30 | 0.77 | -0.087 |  |
| S1 SMS quality | 4.2 (0.6) | 4.0 (0.8) | 1.04 | 0.31 | 0.299 |  |
| S2 SMS quality | 3.8 (0.7) | 3.7 (1.2) | 0.60 | 0.55 | 0.173 |  |
| B. PVT | | | Effect | F | p | Partial $\eta^2$ |
| S1 PVT (ms) | 286 (24) | 298 (28) | Session | 1.32 | 0.26 | 0.028 |
| S2 PVT (ms) | 286 (24) | 293 (23) | Group | 2.11 | 0.15 | 0.044 |
|  |  |  | S x G | 1.61 | 0.21 | 0.034 |
| C. SSS | | | Effect | F | p | Partial $\eta^2$ |
| S1 SSS | 2.1 (0.9) | 2.3 (1.0) | Session | 0.02 | 0.89 | 0.000 |
| S2 SSS | 2.3 (0.8) | 2.2 (1.2) | Group | 0.07 | 0.79 | 0.002 |
|  |  |  | S x G | 0.98 | 0.33 | 0.021 |

#### 2.1.1 Sample size

As the specific effect of M1 stimulation on the integration of new information into a compatible cognitive-motor schema has not previously been investigated, a literature-based a priori estimation of the hypothesized effect size is not possible. Nonetheless, given the identical task and analogous experimental design, we based our sample size calculation on our previous work investigating the influence of compatible vs. incompatible schema on motor memory integration (King et al., 2019). The primary effect of interest in this earlier work was the group main effect in the response time (RT) for novel transitions in the Session 2 training phase, as assessed with a 2 (compatible vs. incompatible) x 20 (practice blocks) ANOVA (F(1,36)=5.29, p=0.027, Ƞp^2^=0.128, Cohen’s F=0.383). Corresponding power analysis (Effect size f=0.383, tails=2; alpha=0.05, Power=0.80, correlation among repeated measures=0.72) via the software G*Power (Faul et al., 2007) resulted in an estimated 21 subjects per group. We aimed to recruit approximately 25 subjects per group to account for participant attrition.

#### 2.1.2 Data exclusions

Out of the 51 volunteers who participated in this study (25 in the STIM group and 26 in the SHAM group), three participants were excluded from all analyses (one participant in the STIM group and two in the SHAM group). Two (one in each group) were excluded for non-compliance with experimental instructions (i.e., one for failing to maintain a regular sleep schedule and the other for failing to comply to task instructions) and another one (SHAM group) for failing to perform the correct motor sequence in Session 1 (percentage of correct sequence transitions during Session 1 greater than 3 standard deviations below the mean of all subjects in the group). Consequently, 48 participants were included in the analyses of the current study (24 participants in the STIM group and 24 participants in the SHAM group).

### 2.2 Motor tasks

The motor task was analogous to that employed in our earlier research (King et al., 2019; Reverberi et al., 2023). Participants performed either a sequential or a pseudo-random variant of the serial reaction time task (SRTT), coded and implemented in MATLAB (The MathWorks Inc., 2019) using Psychophysics Toolbox version 3 (Kleiner et al., 2007). Participants performed the task while seated at a desk and with their hands positioned under a small tray to prevent them from seeing their fingers.

During the motor task, eight squares, corresponding spatially to the 8 fingers used to perform the task (no thumbs were used) were shown on the screen. The outline of the squares was colored red and green, indicating periods of rest or practice, respectively. During practice, the green squares were consecutively filled with cues (full green square) at the different spatial locations on the computer screen, and participants were instructed to press the key corresponding to the cue with the corresponding finger as quickly and accurately as possible. In the sequential version of the task, the cues followed a deterministic 8-element sequence. For this version of the task, participants were explicitly told that the order of the key presses would follow a sequential repetitive pattern, but were not given any additional information regarding the length or structure of the sequence. In the pseudo-random version of the task, each of the eight keys were pressed once in every run of 8 consecutive key presses, ensuring that the key distribution was the same as in the sequential task. Each practice block contained 64 key presses (corresponding to 8 repetitions of the 8-element sequence or 8 consecutive random patterns of 8 key presses) and was followed by a 15-second rest period. The pressed key and the timing of each key press was continuously recorded during the task.

### 2.3 Experimental design

Participants were invited for two experimental sessions that took place approximately 24 hours apart (see design in Fig. 1) and were scheduled during the day between 9AM and 6.30PM. Participants were instructed to begin following a constant sleep/wake schedule (according to their own rhythm, minimum 7h of sleep per night and with a bed time no later than 1.00 AM) 3 nights prior to the first experimental session. Compliance with this sleep schedule was assessed via self-reported sleep diaries, and additionally via a wrist-mounted actigraphy device (ActiGraph wGT3X-BT, Pensacola, FL) for the night between the two experimental sessions. Sleep quality and quantity for the night preceding each experimental session were assessed with the St. Mary’s sleep questionnaire (Ellis et al., 1981). Prior to each experimental session, the psychomotor vigilance task (Dinges & Powell, 1985) and the Stanford Sleepiness Scale (Hoddes, 1972) were also administered to provide objective and subjective assessments of vigilance, respectively.

**Figure 1:**
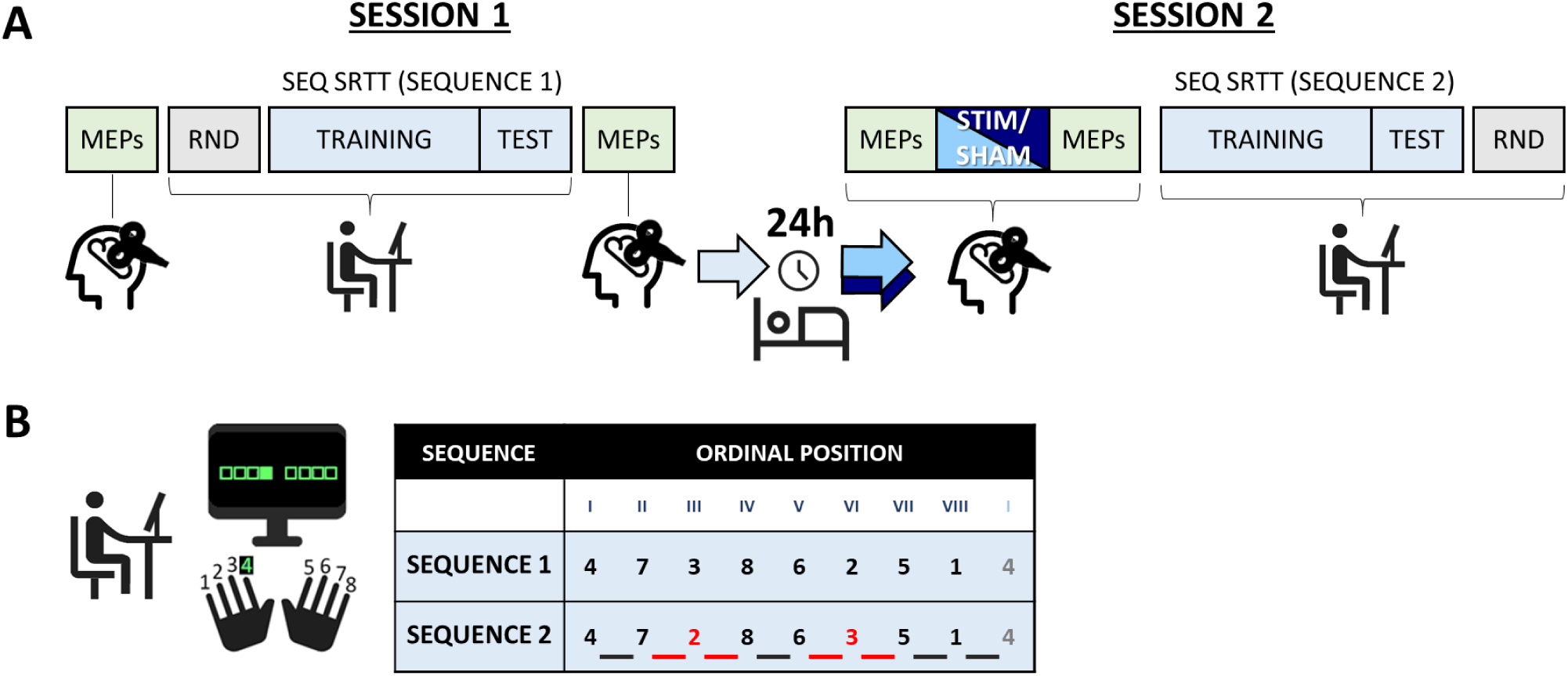
**A.** Experimental design. Participants performed two sessions, approximately 24 hours apart. In Session 1, after TMS baseline measures (determination of hotspot, rMT and pre-learning MEP measurements), participants performed 4 blocks of pseudo-random SRTT (RND), followed by learning the sequential SRTT (SEQ: 20 training blocks, 4 test blocks). MEPs were then collected post-learning. In Session 2, after pre-stimulation MEPs were measured, participants received either sham (SHAM group) or active (STIM group) stimulation of the hotspot. Post-stimulation MEPs were then measured. Next, participants performed the sequential SRTT (20 training and 4 test blocks). The session ended with 4 blocks of the pseudo-random SRTT. **B.** During the sequential SRTT, in Session 1 participants learned Sequence 1 (corresponding to keys 4-7-3-8-6-2-5-1, in which 1 through 8 are the left pinky to the right pinky fingers, excluding the thumbs). In Session 2, they learned a second motor sequence (Sequence 2, 4-7-2-8-6-3-5-1). The ordinal structure of Sequence 2 was highly compatible with that of Sequence 1, as only two keys (highlighted in red) were presented in new ordinal positions (roman numerals in the table). This manipulation allowed us to test for the causal role of M1 in the retention of previously learned transitions (examining the 4 transitions previously learned in Session 1, underlined in black) and in the integration of new movement transitions (examining the 4 novel transitions newly presented in Session 2, underlined in red). Icons used in this figure are adapted from Google Material Symbols (Apache License, version 2.0).

In Session 1 on Day 1, after baseline TMS measurements (i.e., search of hotspot, resting motor threshold (rMT) and pre-learning Motor Evoked Potentials (MEPs) measurements, see Brain stimulation section below), participants performed 4 blocks of the pseudorandom SRTT variant to assess baseline motor performance. These were followed by 20 blocks of the sequential SRTT during which participants learned a bimanual motor sequence (Sequence 1; 4-7-3-8-6-2-5-1, where numbers 1 through 8 correspond to the left pinky to the right pinky fingers, thumbs excluded; Training in Fig. 1). The training blocks were followed by a “test” run of 4 practice blocks. Prior to the test run, participants were allowed approximately 1 minute of rest to assess end-of-training performance following the dissipation of mental and physical fatigue (Pan & Rickard, 2015). MEPs were collected again after completion of the motor task to assess the effect of learning on corticospinal excitability. Participants then received instructions regarding actigraphy device wear and maintenance of a regular sleep schedule, and returned home to follow their habitual daily schedule. In Session 2 on Day 2, after baseline TMS measurements (i.e., search of hotspot, rMT determination, and pre-stimulation MEP measurements) participants received either sham stimulation (SHAM group) or continuous theta burst stimulation to the left M1 (STIM group; duration stimulation/sham sessions: 40 seconds, see Brain stimulation procedures). Several measures were taken to prevent participant detection of group assignment. Firstly, participants were not informed of the presence of active and sham stimulation groups. Secondly, for both groups, the experimenter informed participants that individuals experience the TBS stimulation differently, with experiences ranging between barely detectable to uncomfortable. Finally, in all sessions the experimenter had the TMS coils positioned on a table behind the participant and motioned as if to switch stimulation coil between MEP collection and active/sham TBS (the coil was truly switched only in the SHAM group). Five minutes of rest were offered to the participants after the end of the stimulation/sham sessions. After rest, post-stimulation MEPs were collected to measure the effect of cTBS/sham stimulation on corticospinal excitability. Participants then learned a novel motor sequence (Sequence 2, 20 blocks of training and 4 blocks of test after a short rest interval), which was designed as in our previous research to be highly compatible with Sequence 1 in terms of its ordinal structure (King et al., 2019; Reverberi et al., 2023). Specifically, 75% of the elements in Sequence 2 (4-7-2-8-6-3-5-1) were in the same ordinal positions as in Sequence 1 (i.e., only keys 2 and 3 were in new ordinal positions; see Fig. 1). With this manipulation, 4 of the movement transitions in Sequence 2 were already learned in the previous session (*learned* transitions: 4-7, 8-6, 5-1, 1-4), while the other 4 were not previously learned (*novel* transitions: 7-2, 2-8, 6-3, 3-5). After sequential SRTT performance, participants performed 4 blocks of pseudorandom SRTT at the end of Session 2.

### 2.4 Brain stimulation

TMS stimulation was delivered using a DuoMAG XT-100 repetitive TMS (rTMS) stimulator (DEYMED Diagnostics s.r.o., Hronov, Czech Republic) and online spatial monitoring of the coil position was performed using neuronavigation (BrainSight, Rogue Research Inc, Montreal, Quebec, CA). During each experimental session, prior to the execution of the motor task (in Session 1) or to the stimulation session (in Session 2), the optimal spot for left M1 stimulation (hotspot) was defined as the location over the scalp where single TMS pulses consistently produced the largest-amplitude MEPs in the right first dorsal interosseous (FDI) muscle, as measured with electromyography. The resting motor threshold (rMT) of each participant was defined as the minimal stimulation intensity at the hotspot required to reach MEPs of at least 50µV peak-to-peak amplitude in the right FDI on at least 5 out of 10 consecutive trials (Rossini et al., 2015). During each session, corticospinal excitability was also assessed at 2 timepoints, i.e., preceding and following motor task execution in Session 1, and preceding and following cTBS/sham stimulation administration in Session 2. To do so, 21 MEPs (the first MEP is discarded from the analyses, see section 2.5.2) were recorded from the hotspot as resulting from single-pulse stimulations delivered at intensity of 120% of the rMT.

During Session 2, after the baseline TMS measurements described above but before SRTT performance (see Fig. 1), participants received either active inhibitory (STIM group) or sham stimulation (SHAM group) of the left M1 hotspot. The target region was selected based on our previous work showing left M1 involvement in the integration of novel material into an acquired cognitive-motor schema (Reverberi et al., 2025). Participants in the STIM group received continuous Theta Burst Stimulation (cTBS), a protocol which was shown to result in the suppression of corticospinal excitability (Huang et al., 2005) and that consisted of three pulses delivered at 50Hz frequency and repeated every 200ms for 40s, for a total of 600 pulses, delivered at an intensity of 80% of the rMT (Derosiere et al., 2017a; Derosiere et al., 2017b). Participants in the SHAM group received sham stimulation, which consisted of the same stimulation pattern described above but delivered through a coil fitted with a spacer (3D-printed, plastic, 33mm thickness) to prevent any active stimulation. As active cTBS effects are known to outlast the stimulation protocol by 30 to 60 min (Huang et al., 2005), they therefore overlapped with SRTT performance (the mean time elapsed between collection of MEPs pre-cTBS/sham stimulation and SRTT performance was of 12±1min).

### 2.5 Statistical analyses

Statistical analyses were performed on IBM SPSS Statistics for Windows, version 29 (IBM Corp, 2021). For all repeated-measures ANOVAs detailed below, Greenhouse-Geisser corrections for ε≤0.75, or Huynh-Feldt corrections for ε>0.75, were applied in the event of violation of sphericity (J. P. Verma, 2015).

#### 2.5.1 Behavioral data

The primary outcome variable for the motor tasks were: 1) response time (RT), i.e. time between cue presentation and participant key press, and 2) accuracy, i.e., correctness of response. Individual trials were excluded from the analyses when the measured RT was greater than 3 standard deviations above or below the participant’s mean response time for that block. This led to the exclusion of an average of 1.58% of trials across the two experimental sessions and groups, which is in line with what was observed in our previous study employing the same motor task (Reverberi et al., 2023). The mean RT for correct movement transitions (i.e., both response n and n-1 were correct) and the percentage of correct transitions were computed for each block of practice on the sequential SRT task. For both measures, averaging was done across all transitions (Sessions 1 and 2) as well as on learned and novel transitions separately (Session 2 only) as in our previous research (King et al., 2019; Reverberi et al., 2023).

To test the effect of cTBS versus sham stimulation of the left M1 on Sequence 2 performance, we performed separate repeated-measures ANOVAs on RT and accuracy measures (training and test runs) with the between-subject factor *group* (STIM/SHAM) and the within-subject factor *block*. This analysis was performed on all movement transitions, as well as on two subsets of movement transitions: a) novel transitions (investigating the effect of M1 stimulation on integration processes) and b) learned transitions (investigating the effect of M1 stimulation on memory retention processes). Additionally, repeated-measures ANOVAs with the between-subject factor *group* (STIM/SHAM) and the within-subject factor *transition type* (novel/learned) were performed on RT and accuracy measures from the Session 2 post-training test run to compare performance related to memory integration and retention between the two groups after extensive task practice.

Negative control analyses were performed on vigilance scores (i.e., mean reaction time on the PVT task and Stanford Sleepiness Scale, assessed with *session* x *group* ANOVA), sleep quantity and quality (St. Mary’s sleep questionnaire (Ellis et al., 1981), independent-samples t-tests), and inclusion questionnaire scores (assessed with independent-samples t-tests, see section Participants for details).

Control analyses were also performed on the pseudo-random SRTT data separately for the two experimental sessions (RT and accuracy, separate *group* x *block* ANOVAs) to test whether baseline motor execution differed between experimental groups (Session 1) and whether any potential group differences observed on the sequential SRTT performance could be attributed to general differences in motor execution at the end of the experiment (Session 2). These results are presented in the Supplementary material.

Finally, exploratory analyses were performed within the STIM group to compare SRTT performance between TMS responders and non-responders, defined based on the amplitude of the cTBS-induced changes in MEP amplitude during Session 2. Specifically, participants displaying an MEP suppression (i.e. decrease in MEP amplitude from pre-to post-cTBS) were considered responders (n=13), while participants displaying an MEP increase were considered non-responders (n=11). Performance was compared between responders and non-responders with repeated-measures ANOVAs analogous to those described above. Corresponding results are presented in the Supplementary material.

#### 2.5.2 Corticospinal excitability

Twenty-one MEPs were collected at two different timepoints in both sessions to assess (i) learning-related changes in corticospinal excitability (MEPs collected pre- and post-learning in Session 1) and (ii) stimulation-induced changes in corticospinal excitability (MEPs collected pre- and post-stimulation in Session 2). Due to experimental error or participant discomfort, data of 3 participants (2 in the SHAM group and 1 in the STIM group) included fewer MEPs (with a minimum of 11 MEPs per timepoint). To correct DC offset, the mean baseline amplitude (in the 50ms pre-TMS pulse) was first subtracted from each MEP signal. For all participants, the first MEP was then excluded from analyses as its amplitude is usually higher than that of subsequent MEPs due to reflex or startle responses. Other MEPs were excluded from analyses if their amplitude was lower than 50µV (a total of 2.50% of all available MEPs were excluded for this reason) and in case of high background muscle activity, i.e., the root mean square of the signal exceeded 20 μV during the 50ms pre-TMS pulse (Cuypers et al., 2020; a total of 0.11% of all available MEPs was excluded for this reason). Following these data exclusions, one participant in the SHAM group was entirely excluded from Session 1 MEP analyses as too few MEPs remained for the second timepoint (<5 MEPs; Osnabruegge et al., 2023). An average of 19 MEPs per timepoint, per participant, remained for analyses (range across all participants, sessions and timepoints: 7-20). For the remaining motor-evoked potentials, peak-to-peak amplitude was calculated in the 50ms post-TMS pulse and then averaged per timepoint and per participant. In Session 1, a paired t-test (pre-versus post-SRTT) was performed to test the effect of motor learning on corticospinal excitability. In Session 2, a repeated-measures ANOVA was performed with the between-subject factor *group* (SHAM/STIM) and the within-subject factor *timepoint* (pre-/post-stimulation) to test for the effect of the stimulation protocol on corticospinal excitability. Additionally, MEP amplitude changes (post-pre) were calculated for each session and participant. Exploratory analyses were performed to test for a relationship between the change in MEP amplitude and performance on the sequential SRTT (response time and accuracy for all, learned, and novel transitions).

## 3. Results

### 3.1 Participant characteristics

Participant characteristics, vigilance at time of testing, as well as sleep quantity and quality for the nights preceding the experimental sessions are presented in Table 1, along with the results of the corresponding statistical analyses comparing metrics between experimental groups. Performance on the random SRTT task, measured at the start of Session 1 and the end of Session 2, is depicted in Fig. 2A and corresponding statistical analyses are reported in Table S1. In brief, the two experimental groups did not significantly differ in any of the reported measures.

**Figure 2:**
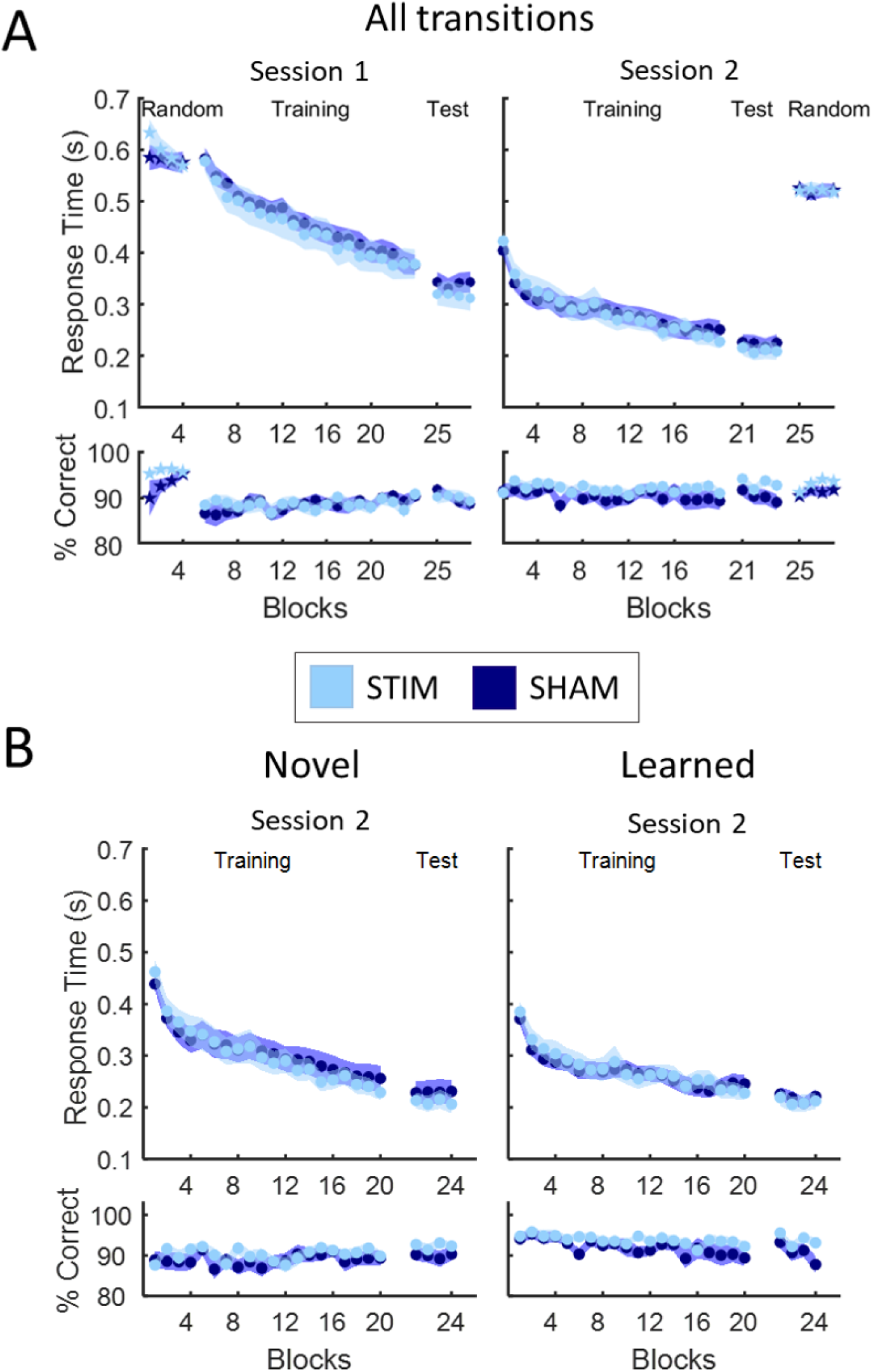
SRTT performance. Mean response time and accuracy for correct keypresses (random SRTT; star markers) or correct movement transitions (sequential SRTT; circle markers), averaged per block of practice, are shown separately for the STIM group (light blue) and SHAM group (dark blue). Performance averaged across all movement transitions (A) or separately across learned and novel transitions (B, Session 2 only). Shaded areas represent the standard error of the mean (SEM).

### 3.2 Behavioral results

Sequential SRTT performance on all transitions is depicted in Fig. 2A and the output of the corresponding statistical analyses is presented in Table 2A.

**Table 2:**
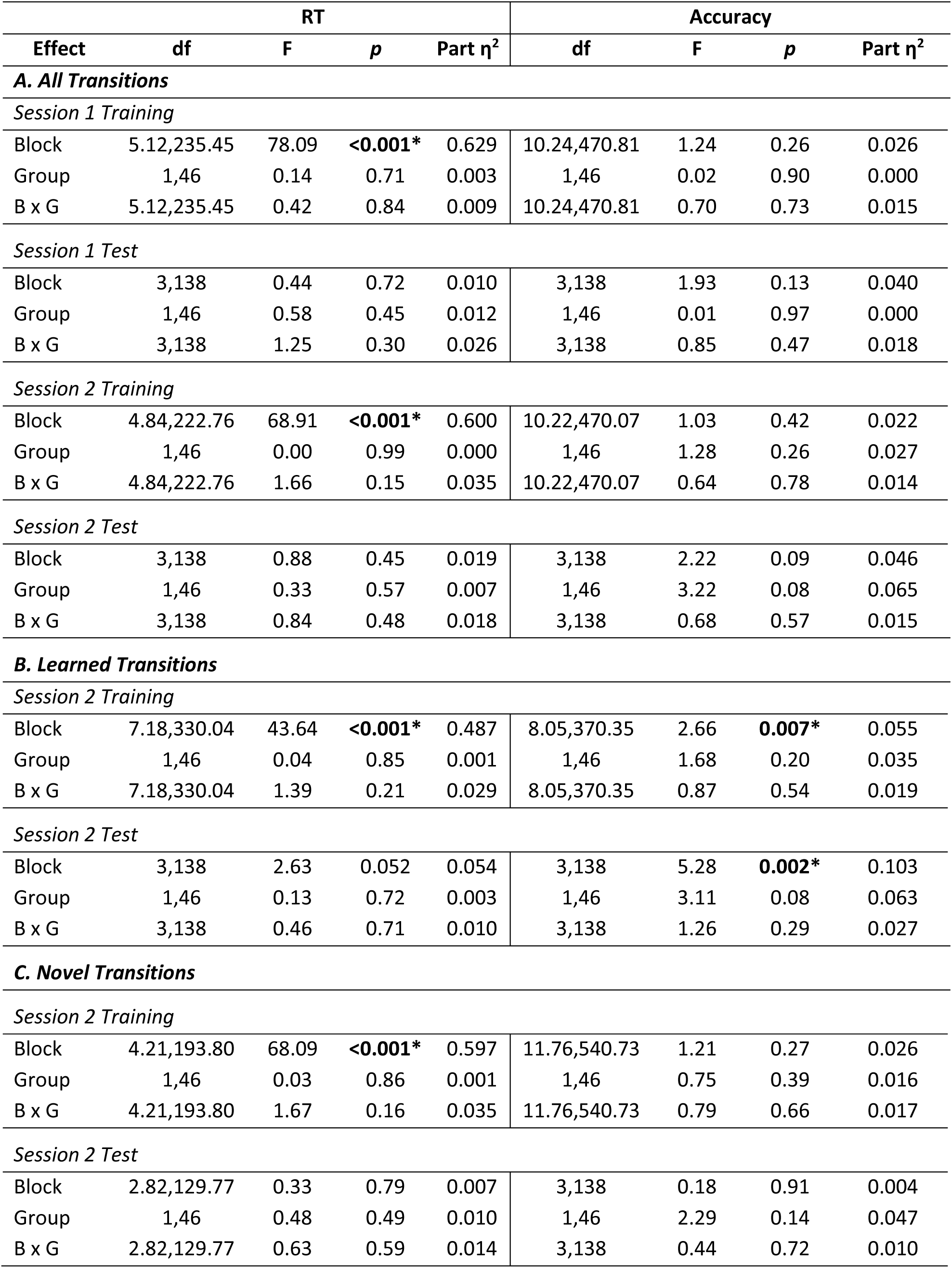
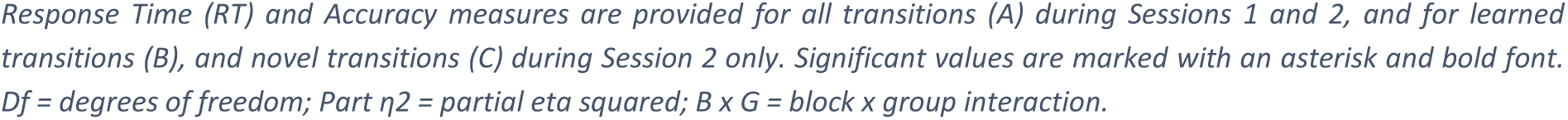
Results of the statistical analyses on the sequential SRTT performance.

| RT |  |  |  |  | Accuracy |  |  |  |
| --- | --- | --- | --- | --- | --- | --- | --- | --- |
| Effect | df | F | p | Part $\eta^2$ | df | F | p | Part $\eta^2$ |
| **A. All Transitions** |  |  |  |  |  |  |  |  |
| *Session 1 Training* |  |  |  |  |  |  |  |  |
| Block | 5.12,235.45 | 78.09 | **<0.001\*** | 0.629 | 10.24,470.81 | 1.24 | 0.26 | 0.026 |
| Group | 1,46 | 0.14 | 0.71 | 0.003 | 1,46 | 0.02 | 0.90 | 0.000 |
| B x G | 5.12,235.45 | 0.42 | 0.84 | 0.009 | 10.24,470.81 | 0.70 | 0.73 | 0.015 |
| *Session 1 Test* |  |  |  |  |  |  |  |  |
| Block | 3,138 | 0.44 | 0.72 | 0.010 | 3,138 | 1.93 | 0.13 | 0.040 |
| Group | 1,46 | 0.58 | 0.45 | 0.012 | 1,46 | 0.01 | 0.97 | 0.000 |
| B x G | 3,138 | 1.25 | 0.30 | 0.026 | 3,138 | 0.85 | 0.47 | 0.018 |
| *Session 2 Training* |  |  |  |  |  |  |  |  |
| Block | 4.84,222.76 | 68.91 | **<0.001\*** | 0.600 | 10.22,470.07 | 1.03 | 0.42 | 0.022 |
| Group | 1,46 | 0.00 | 0.99 | 0.000 | 1,46 | 1.28 | 0.26 | 0.027 |
| B x G | 4.84,222.76 | 1.66 | 0.15 | 0.035 | 10.22,470.07 | 0.64 | 0.78 | 0.014 |
| *Session 2 Test* |  |  |  |  |  |  |  |  |
| Block | 3,138 | 0.88 | 0.45 | 0.019 | 3,138 | 2.22 | 0.09 | 0.046 |
| Group | 1,46 | 0.33 | 0.57 | 0.007 | 1,46 | 3.22 | 0.08 | 0.065 |
| B x G | 3,138 | 0.84 | 0.48 | 0.018 | 3,138 | 0.68 | 0.57 | 0.015 |
| **B. Learned Transitions** |  |  |  |  |  |  |  |  |
| *Session 2 Training* |  |  |  |  |  |  |  |  |
| Block | 7.18,330.04 | 43.64 | **<0.001\*** | 0.487 | 8.05,370.35 | 2.66 | **0.007\*** | 0.055 |
| Group | 1,46 | 0.04 | 0.85 | 0.001 | 1,46 | 1.68 | 0.20 | 0.035 |
| B x G | 7.18,330.04 | 1.39 | 0.21 | 0.029 | 8.05,370.35 | 0.87 | 0.54 | 0.019 |
| *Session 2 Test* |  |  |  |  |  |  |  |  |
| Block | 3,138 | 2.63 | 0.052 | 0.054 | 3,138 | 5.28 | **0.002\*** | 0.103 |
| Group | 1,46 | 0.13 | 0.72 | 0.003 | 1,46 | 3.11 | 0.08 | 0.063 |
| B x G | 3,138 | 0.46 | 0.71 | 0.010 | 3,138 | 1.26 | 0.29 | 0.027 |
| **C. Novel Transitions** |  |  |  |  |  |  |  |  |
| *Session 2 Training* |  |  |  |  |  |  |  |  |
| Block | 4.21,193.80 | 68.09 | **<0.001\*** | 0.597 | 11.76,540.73 | 1.21 | 0.27 | 0.026 |
| Group | 1,46 | 0.03 | 0.86 | 0.001 | 1,46 | 0.75 | 0.39 | 0.016 |
| B x G | 4.21,193.80 | 1.67 | 0.16 | 0.035 | 11.76,540.73 | 0.79 | 0.66 | 0.017 |
| *Session 2 Test* |  |  |  |  |  |  |  |  |
| Block | 2.82,129.77 | 0.33 | 0.79 | 0.007 | 3,138 | 0.18 | 0.91 | 0.004 |
| Group | 1,46 | 0.48 | 0.49 | 0.010 | 1,46 | 2.29 | 0.14 | 0.047 |
| B x G | 2.82,129.77 | 0.63 | 0.59 | 0.014 | 3,138 | 0.44 | 0.72 | 0.010 |

Response Time (RT) and Accuracy measures are provided for all transitions (A) during Sessions 1 and 2, and for learned transitions (B), and novel transitions (C) during Session 2 only. Significant values are marked with an asterisk and bold font. $Df$ = degrees of freedom; $\text{Part } \eta^2$ = partial eta squared; $B \times G$ = block × group interaction.

During training in Session 1, participants of both experimental groups successfully and similarly learned the motor sequence, as evidence by the significant effect of *block* on RT reflecting progressively faster performance, and the lack of *group* effect or *block* x *group* interaction. During the post-training test, the lack of any significant effects on RT indicates that performance reached similar plateau levels for both groups. Accuracy remained high and stable throughout all of Session 1, with no difference between groups.

Analyses of all transitions of Session 2 revealed a significant effect of *block* on RT during training, but not during test, and no significant *group* effect or *block* x *group* interactions, suggesting that participants in both groups similarly learned the second motor sequence. Accuracy on all transitions remained high and stable. These analyses were repeated on two subsets of movements transitions, i.e. the learned transitions (those that had been previously learned in Session 1) and the novel transitions (that were introduced during Session 2, Fig. 2B). These analyses, reported in Table 2B-C, revealed analogous results as those observed for all transitions, i.e., no between-group differences in either RT or accuracy.

Finally, repeated-measures ANOVAs with the between-subject factor *group* and the within-subject factor *transition type* (novel vs. learned) were performed on the Session 2 test data. The results revealed no group differences in performance between learned and novel transitions after extensive task practice (**response time** *–* main effect *transition type*: *F*(1,46)=0.35, *p*=0.56, η^2^=0.007; main effect *group*: *F*(1,46)=0.35, *p*=0.56, η^2^=0.008; interaction: *F*(1,46)=0.47, *p*=0.50, η^2^=0.010; **accuracy** – main effect *transition type*: *F*(1,46)=2.75, *p*=0.10, η^2^=0.056; main effect *group*: *F*(1,46)=3.23, *p*=0.08, η^2^=0.066; interaction: *F*(1,46)=0.21, *p*=0.65, η^2^=0.004).

Additionally, we performed exploratory analyses to determine whether Session 2 performance differed between participants classified as cTBS-responders (n=13) and non-responders (n=11, see methods). The results of these analyses are reported in supplementary Table S2 and Figures S1 and S2 and revealed no performance difference between these sub-groups.

Altogether, this pattern of results suggests that cTBS applied to the left M1 prior to learning the new sequence during Session 2 did not influence general motor performance, nor the retention of previously learned motor information or the integration of novel information into a compatible schema.

### 3.3 Motor-evoked potentials

First, we performed an exploratory analysis similar to that performed by Tunovic and colleagues (2014) to examine whether sequence learning modulated cortico-spinal excitability. To do so, we contrasted MEP amplitude from pre-to post-SRTT practice in Session 1. A significant decrease in MEP amplitude was observed across both groups after sequential SRTT practice (t_46_=2.62, *p*=0.01, Cohen’s d=649.81), suggesting that learning of the motor sequence resulted in a decrease in corticospinal excitability (see Fig. 3).

**Figure 3:**
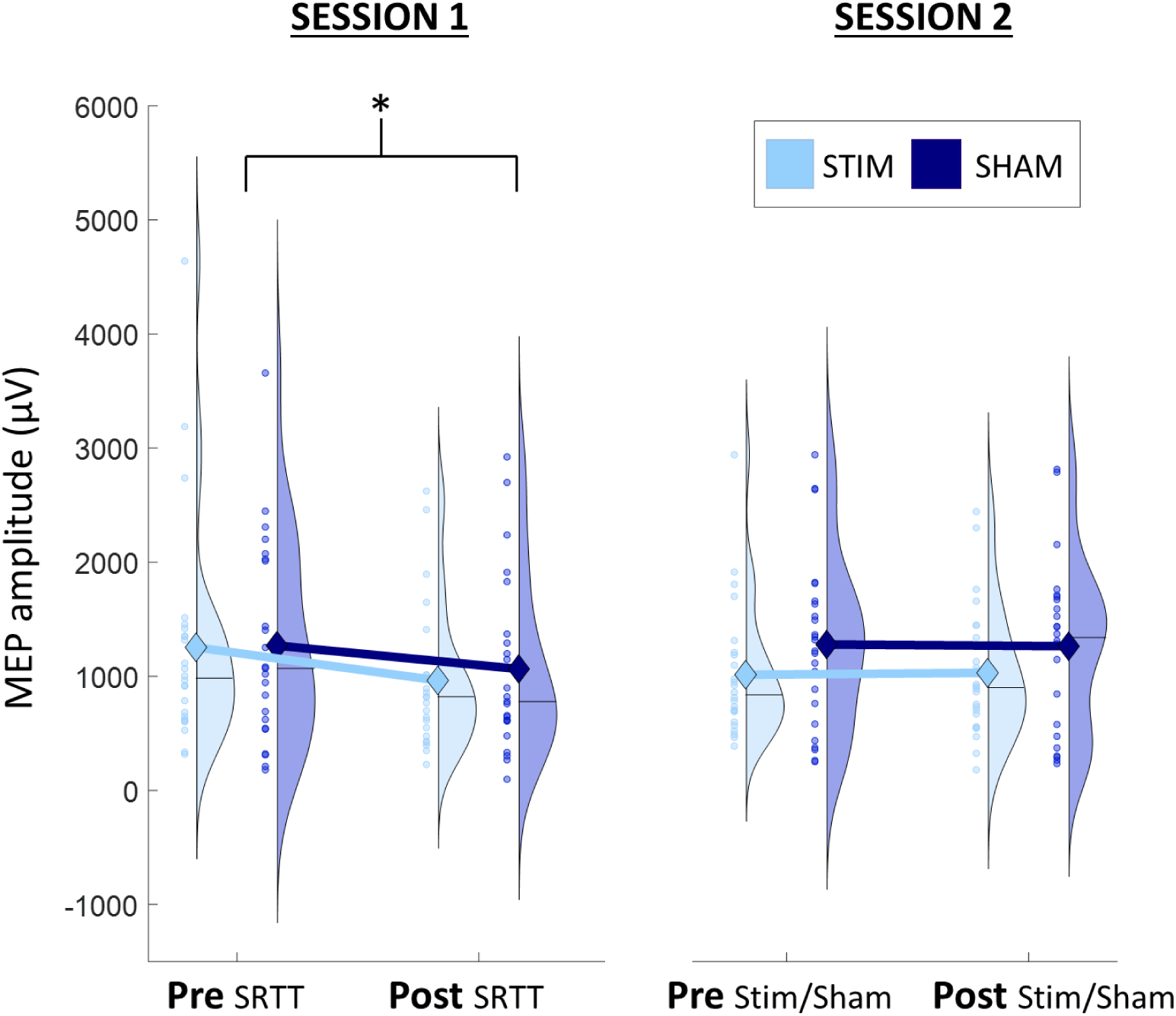
Mean amplitude of motor-evoked potentials, presented separately per timepoint and per group (STIM group: light blue, SHAM group: dark blue). Timepoints are pre- and post-SRTT in Session 1 and pre- and post-cTBS/sham in Session 2. Diamonds indicate the mean and black horizontal bars the medians. Coloured circles represent individual data. The shape of the half-violin depicts the kernel density estimate of the data. Significant effects are indicated with an asterisk.

We then tested whether the cTBS intervention in Session 2 modulated cortico-spinal excitability compared to sham stimulation. MEP amplitude did not significantly change after cTBS/sham intervention (*timepoint* effect, *F*(1,46)=0.00, *p*=0.99, η^2^=0.000) and no significant effect of *group* or *timepoint* x *group* interaction were observed (main effect of *group: F*(1,46)=1.90, *p*=0.18, η^2^=0.040; *timepoint* x *group* interaction: *F*(1,46)=0.06, *p*=0.82, η^2^=0.001). These results suggest that, contrary to our expectations, inhibitory cTBS did not significantly decrease corticospinal excitability in the STIM group (Fig. 3).

### 3.4 Exploratory MEP analyses

#### 3.4.1 Relationship between MEP changes and performance

First, we examined whether the learning-related changes in MEP amplitude observed during Session 1 correlated with sequential SRTT performance across the two experimental groups. There was no significant correlation between MEP change in Session 1 and performance on the 20 training blocks (*RT*: r=-0.23, *p*=0.12; *accuracy*: r=-0.11, *p*=0.46) nor between MEP change and Session 1 online gains computed as the difference between performance in block 1 and average performance on the 4 test blocks (*RT*: r=-0.14, *p*=0.35; *accuracy*: r=-0.11, *p*=0.48).

Second, we examined the relationship between MEP amplitude changes as a function of cTBS or sham stimulation and sequential SRTT performance in Session 2. The results of these analyses are reported in Table 3A-B. No significant correlations were observed within either experimental group and no correlations were significantly different between the two groups.

**Table 3:**
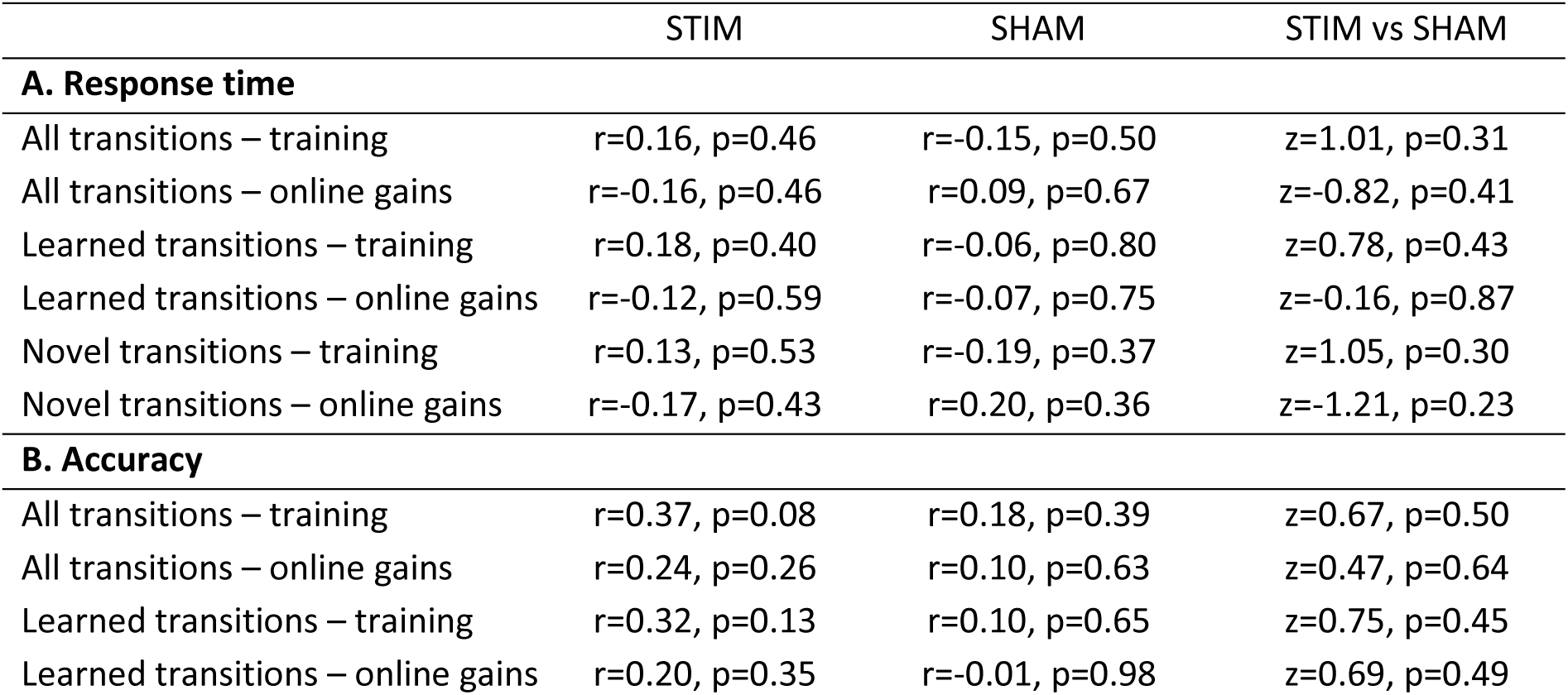
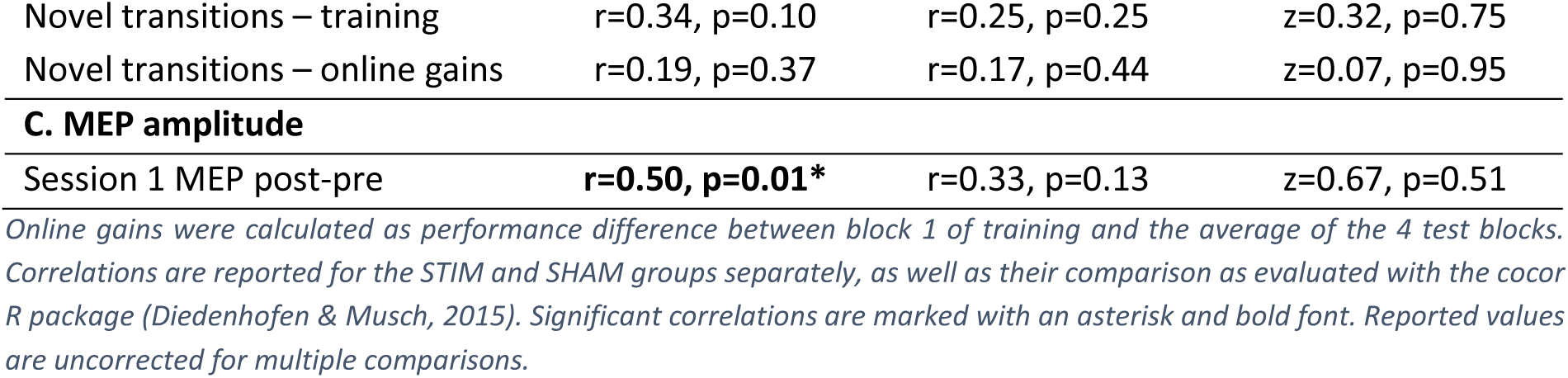
Results of the correlation analyses between Session 2 change in MEP amplitude (post-pre) and: (A) response time in the Session 2 sequential SRTT, (B) accuracy in the Session 2 sequential SRTT, (C) Session 1 change in MEP amplitude.

|  | STIM | SHAM | STIM vs SHAM |
| --- | --- | --- | --- |
| **A. Response time** |  |  |  |
| All transitions – training | $r=0.16$, $p=0.46$ | $r=-0.15$, $p=0.50$ | $z=1.01$, $p=0.31$ |
| All transitions – online gains | $r=-0.16$, $p=0.46$ | $r=0.09$, $p=0.67$ | $z=-0.82$, $p=0.41$ |
| Learned transitions – training | $r=0.18$, $p=0.40$ | $r=-0.06$, $p=0.80$ | $z=0.78$, $p=0.43$ |
| Learned transitions – online gains | $r=-0.12$, $p=0.59$ | $r=-0.07$, $p=0.75$ | $z=-0.16$, $p=0.87$ |
| Novel transitions – training | $r=0.13$, $p=0.53$ | $r=-0.19$, $p=0.37$ | $z=1.05$, $p=0.30$ |
| Novel transitions – online gains | $r=-0.17$, $p=0.43$ | $r=0.20$, $p=0.36$ | $z=-1.21$, $p=0.23$ |
| **B. Accuracy** |  |  |  |
| All transitions – training | $r=0.37$, $p=0.08$ | $r=0.18$, $p=0.39$ | $z=0.67$, $p=0.50$ |
| All transitions – online gains | $r=0.24$, $p=0.26$ | $r=0.10$, $p=0.63$ | $z=0.47$, $p=0.64$ |
| Learned transitions – training | $r=0.32$, $p=0.13$ | $r=0.10$, $p=0.65$ | $z=0.75$, $p=0.45$ |
| Learned transitions – online gains | $r=0.20$, $p=0.35$ | $r=-0.01$, $p=0.98$ | $z=0.69$, $p=0.49$ |
| Novel transitions – training | $r=0.34, p=0.10$ | $r=0.25, p=0.25$ | $z=0.32, p=0.75$ |
| Novel transitions – online gains | $r=0.19, p=0.37$ | $r=0.17, p=0.44$ | $z=0.07, p=0.95$ |
| C. MEP amplitude |  |  |  |
| Session 1 MEP post-pre | **$r=0.50, p=0.01^*$** | $r=0.33, p=0.13$ | $z=0.67, p=0.51$ |

Online gains were calculated as performance difference between block 1 of training and the average of the 4 test blocks. Correlations are reported for the STIM and SHAM groups separately, as well as their comparison as evaluated with the *cocor* R package (Diedenhofen & Musch, 2015). Significant correlations are marked with an asterisk and bold font. Reported values are uncorrected for multiple comparisons.

#### 3.4.2 Relationship between MEP changes across the two experimental sessions

Finally, to examine whether participants who were more susceptible to undergo learning-induced changes in corticospinal excitability also displayed stronger stimulation-induced MEP modulation, we tested for a potential relationship between changes in MEP amplitude across the two experimental sessions. Results show that a greater learning-related decrease in MEP amplitude in Session 1 was correlated with a greater stimulation/sham-induced decrease in MEP amplitude in Session 2. This correlation was significant on the entire sample (r=0.41, *p*=0.004), as well as within the STIM group, but not within the SHAM group (see Table 3C, Fig. 4A), although correlations were not significantly different between the STIM and SHAM groups. Interestingly, this correlation was particularly pronounced within STIM group participants classified as responders (Fig. 4B), i.e., those who displayed MEP suppression post-cTBS intervention (n=13, r=0.80, *p*=0.001) while it was non-significant in non-responders (n=11, r=-0.001, *p*≅1.00). [Note that similar control analyses were performed on SHAM participants labelled as responders and non-responders with the same criteria as for the STIM participants (see above) and that no significant correlations were detected for these SHAM responders (n=13, r=0.17, *p*=0.58) or SHAM non-responders (n=10, r=0.36, *p*=0.31) sub-groups].

**Figure 4:**
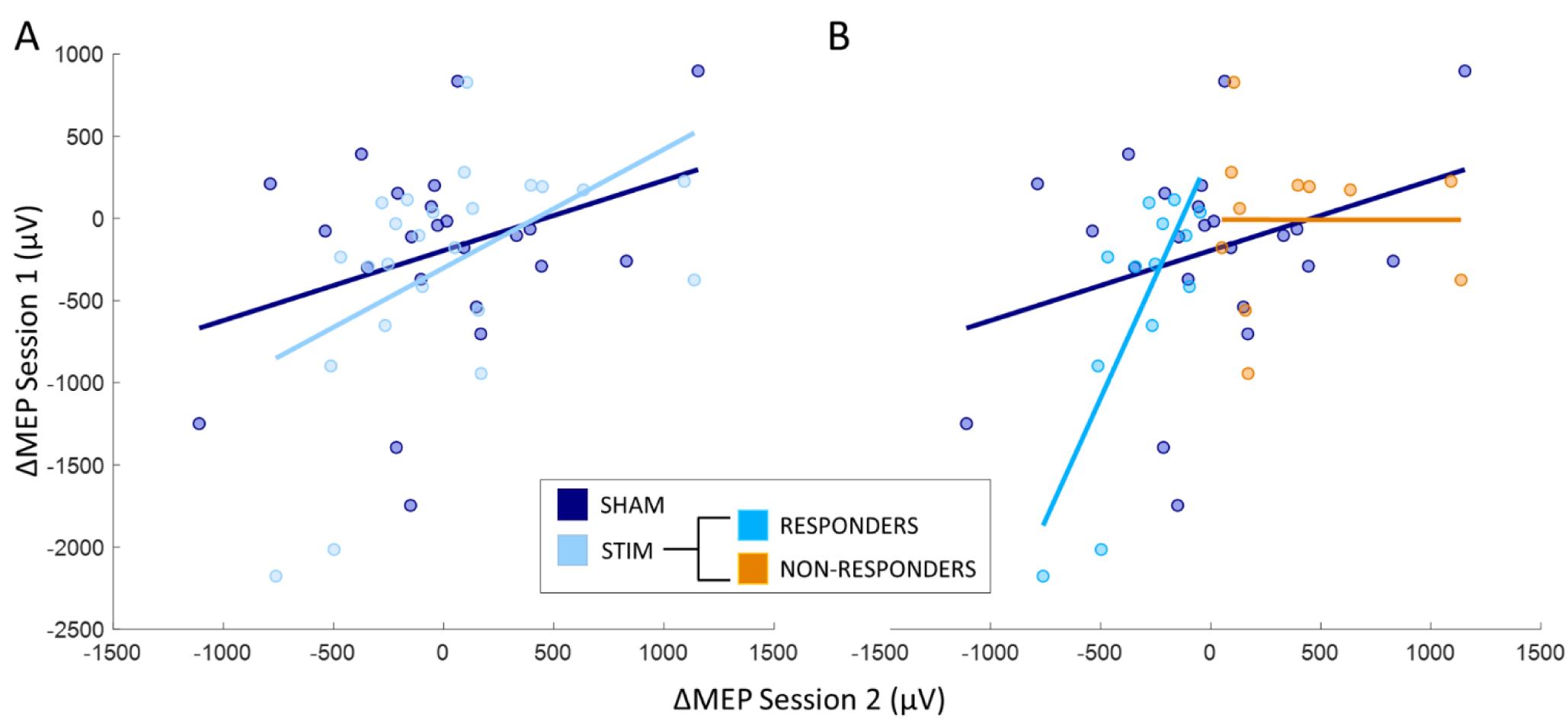
Correlation between changes in amplitude of motor-evoked potentials during the two experimental sessions, presented separately per group (A; STIM group: light blue, SHAM group: dark blue) and per subgroup (B; STIM group responders: blue, STIM group non-responders: orange). Coloured circles represent individual data, lines represent the linear trend.

## 4. Discussion

In the current experiment, we investigated whether inhibitory TMS applied to the left primary motor cortex prior to task performance influenced the integration of a novel motor sequence into a previously acquired cognitive-motor schema. To do so, participants first learned a bimanual motor sequence from which a cognitive-motor schema is thought to be extracted (King et al., 2019). Approximately 24 hours later, participants learned a novel sequence that included both novel movement transitions as well as movement transitions that were learned on Day 1. Importantly, the novel sequence learned on Day 2 was compatible with the ordinal-based cognitive motor schema learned on Day 1. Inhibitory M1 stimulation was delivered prior to learning on Day 2 to examine the effect of M1 cTBS on (1) the retention of previously learned movements and (2) the learning of new movements that are performed in a context that is compatible with the previous schema. Contrary to our expectations, our results did not show any effect of M1 stimulation on motor performance and therefore do not provide evidence for a causal role of M1 in schema-mediated motor learning.

Our results suggest that M1 cTBS did not influence performance on either the previously learned or the novel movements performed on Day 2 of the experiment. These findings are partially in line with prior literature as there is evidence to suggest that inhibitory stimulation applied to the M1 contralateral to the hand used for task practice differently affects sequential motor performance depending on the motor learning stage (e.g., novel vs. well-learned movements). For example, earlier work has shown that disruptive 1Hz repetitive TMS delivered to M1 24 hours after learning a unimanual motor sequence task did not affect finger-tapping performance at an immediate post-stimulation retest (Hotermans et al., 2008). Our results on the learned transitions are in line with these earlier observations, which suggest that inhibitory M1 stimulation does not affect the execution of consolidated motor memory traces. In contrast, our results on the novel transitions contradict a series of experiments showing that inhibitory stimulation applied prior to learning new unimanual sequences of movements significantly impairs motor performance. Specifically, it was demonstrated that inhibitory cTBS administration to M1 before learning of a new unimanual probabilistic sequence task impaired performance during early training blocks (Rosenthal et al., 2009). Similarly, M1 cTBS applied prior to implicit probabilistic learning of a unimanual sequence of keypresses hindered learning as measured by the response time advantage for probable versus improbable targets (Wilkinson et al., 2010). A similar experiment demonstrated that this cTBS-induced impairment of probabilistic SRT learning also persisted 30min after the end of learning, suggesting that M1 is necessary for both initial learning of new motor sequences as well as for short-term recall (Wilkinson et al., 2015).

The origin of the discrepancy between these earlier studies and the current data is unclear. It is unlikely that stimulation procedure contributed to these discrepancies, as we employed a similar stimulation protocol, with similar timing as in previous research (Rosenthal et al., 2009; Wilkinson et al., 2010, 2015), and certainly within the time-window during which cTBS-induced decrease in M1 excitability has been previously observed (Huang et al., 2005). It is however possible that the differences in task design between the current experiment and previous literature may have contributed to these inconsistencies. First, the majority of the studies discussed above used probabilistic implicit tasks whereas the task employed in this research was explicit and used a deterministic sequence. As there is evidence in the literature suggesting that TMS over M1 may differentially affect implicit and explicit sequence learning (Breton & Robertson, 2017; Tunovic et al., 2014), one could argue that the awareness of the sequential material might have contributed to these differences. Support for this view is however limited to studies in which stimulation was applied after learning in order to affect offline consolidation; a process that was not targeted by stimulation in the current research. Second, it is possible that the complexity of the bimanual task used in our research influenced the effect of stimulation on performance as compared to earlier research described above using simpler, unimanual tasks. This is only partially supported by the existing literature as there is evidence showing that M1 stimulation is more (Gerloff et al., 1998) or less (Clark et al., 2019) efficient to modulate motor performance on complex as compared to simple tasks.

Perhaps more importantly, previous research systematically employed unimanual tasks with stimulation applied contralateral to the hand used to perform the task while the current study used a bimanual task with left M1 stimulation. It is therefore possible that the left M1 stimulation delivered here specifically disrupted performance on the contralateral right hand. To test for this possibility, we performed exploratory analyses in which performance was averaged across fingers of the same hand (see supplementary Fig. S3 and Table S3 for details). Contrary to our expectations, we observed a steeper learning curve in later stages of learning in the right hand, contralateral to the stimulation, following cTBS compared with sham. However, this effect was only transient as it was no longer observed during the post-training test. Our results also indicated that active stimulation – as compared to sham – resulted in higher accuracy during post-training test for novel keys performed with the left hand, ipsilateral to the stimulation. These results thus do not support the hypothesis that left M1 cTBS selectively disrupted right-hand performance.

A last point that is worth putting forward is the possibility that the effect of stimulation was limited to a subset of novel movements. Our earlier research (Reverberi et al., 2025) indeed showed that M1 multivoxel patterns were particularly altered on those sequential elements neighboring the first novel key in the sequence stream. It is therefore possible that the effect of M1 stimulation may be limited to specific sequential movements. Consistent with the single hand analyses presented above, our exploratory analyses contrasting group performance at the level of single sequential elements revealed a marginally significant group difference in accuracy during post-training test, particularly for the second novel key in the sequence (see supplementary Fig. S4 and Tables S4-5). Again, contrary to our hypothesis, this effect was driven by higher, rather than poorer, accuracy post-cTBS. Altogether, the additional exploratory analyses suggest that task properties (e.g., sequence awareness, complexity, bimanual vs. unimanual) are unlikely to explain the lack of disruptive effect observed following cTBS intervention in the current study. We discuss below the possibility that the stimulation protocol failed to induce a decrease in M1 excitability.

The cTBS intervention in the current study failed to elicit a decrease in MEP amplitude, suggesting that the stimulation protocol was not effective in modulating M1 excitability. This is in contrast to previous work showing a decrease of MEP amplitude post-cTBS (Huang et al., 2005); however, other studies have also reported a failure of M1 cTBS to elicit MEP suppression at the group level (Do et al., 2018; Jannati et al., 2017). Inspection of MEP distribution (see supplementary Fig. S1) indicated a normal distribution with a zero median whereby approximately half of the participants displayed cTBS-induced MEP suppression (i.e., a decrease in MEP amplitude from pre-to post-intervention, n=13) while the rest of the sample showed increases in MEP amplitude post-stimulation (n=11). This is in line with a recent large meta-analysis demonstrating that cTBS only induced MEP suppression in approximately 65% of participants, with no evidence for a clear division of the population into cTBS-responders and non-responders (Corp et al., 2020). Our exploratory analyses comparing motor performance between responders (showing a negative change in MEP post-stimulation) and non-responders (showing a positive change in MEP post-stimulation) did not yield any significant differences between these sub-groups, as classified in this study (see supplementary Fig. S2 and Table S2). Although this experiment was not powered and designed to test for such differences, the results of these exploratory analyses suggest no effect of stimulation on motor performance in the cTBS-responders.

There is a variety of experimental design choices – known to influence the effect of cTBS on MEP amplitude – that might have contributed to the null MEP results in the current study. First, while cTBS was shown to induce MEP suppression up to 60 minutes post-stimulation (Huang et al., 2005), meta-analyses suggest that this effect is maximal within a much shorter timeframe (within 5 min post-stimulation, Chung et al., 2016) and no longer significant 10-20 minutes post-stimulation (Corp et al., 2020). In our study, MEPs were measured 5 minutes after the TBS intervention; however, the SRT task was administered on average 12 min after stimulation (range: 10-15 min). We tested the possibility that the time elapsed between the stimulation and the start of sequence practice was related to motor performance in the current study (see Supplementary material). The results of these exploratory correlation analyses did not yield any significant results, which suggests that the time elapsed between the TMS intervention and the task practice did not affect sequential performance, at least within the relatively narrow temporal window probed in the current research. Second, it is possible that the stimulation intensity in the current study failed to induce MEP suppression. It has indeed been shown that single-pulse stimulation intensity affects the detection of post-cTBS MEP suppression, with greater suppression being elicited by higher stimulus intensities such as 150 to 180% of the rMT (Goldsworthy et al., 2016; Vallence et al., 2015). We argue that this is unlikely, as previous research using similar stimulation intensities as in the current study was able to show MEP suppression (e.g., Wilkinson et al., 2015). Another factor that could have influenced our results is the time of day at which stimulation was applied, as the time of the stimulation session was spread between 9am-6.30pm across participants (Sale et al., 2007, but see Corp et al., 2020). To test for this possibility, we performed exploratory correlation analyses between the stimulation session time of day and (i) baseline MEP amplitudes as well as (ii) MEP changes observed during either experimental session. Results of these analyses are presented in the Supplemental information and showed no correlations between time of day and any of the MEP measures. Lastly, additional exploratory analyses presented in the Supplemental information determined that participant age and vigilance, which were previously shown to influence individual response to TMS (Bashir et al., 2014; Conte et al., 2008; Müller-Dahlhaus et al., 2008; Stefan et al., 2004) did not correlate with response to TMS in the current study. It is also worth noting that other factors known to influence TMS responses such as intake of nicotine, alcohol, and other drugs (Kähkönen et al., 2002; Lang et al., 2007; Swayne et al., 2009; Ziemann et al., 2015) were controlled for in the current study. Altogether, the results of these exploratory analyses suggest that it is unlikely that our experimental design or participants characteristics affected cTBS-induced MEP modulation in the current study.

As the MEP measurements did not capture the potential effect of the cTBS intervention, it is reasonable to question whether MEPs represent a valid proxy of corticospinal excitability in the current study. To test for this hypothesis, we examined changes in MEP amplitude from pre-to post-sequential practice in Session 1. In line with previous literature (Tunovic et al., 2014), we observed a significant decrease of MEP amplitude after motor sequence learning, which suggests that the sequence learning induced neural plasticity in M1 that was reflected by changes in MEP amplitude. We additionally observed a significant correlation between changes in MEP amplitude in Session 1 (from pre-to post-sequence learning) and Session 2 (from pre-to post-stim/sham intervention). This correlation only reached significance within the STIM group (although correlations were not significantly different between STIM and SHAM) and was particularly strong within those STIM group participants who responded to the intervention (i.e., those who displayed post-cTBS MEP suppression), but not present in SHAM group “responders” (i.e., displaying post-sham MEP suppression). These correlations suggest that the corticospinal excitability of a subset of participants was more susceptible to be modulated by both learning and stimulation interventions. Additionally, we detected a significant negative correlation between pre-sequence learning (Session 1) MEP amplitude and its change post-learning over the entire sample, as well as between pre-stim/sham (Session 2) MEP measures and MEP amplitude change post-STIM and post-SHAM responders (see supplementary Table S7 for details). Although these correlations are exploratory in nature, their observation across both experimental sessions and both groups of responders suggests that participants with higher baseline cortical excitability displayed stronger MEP suppression irrespective of the intervention (learning, stimulation or sham). These findings are partly in line with previous work demonstrating a relationship between baseline levels of cortical inhibition and individual response to rTMS (Daskalakis et al., 2006). The current data further suggests that individuals with higher baseline MEP tended to regress toward the mean during the post measurement irrespective of the intervention, which might reflect a homeostatic rather than a plasticity process. In line with this interpretation, we did not observe any correlations between baseline MEP measurements and changes in performance. Specifically, we tested whether baseline MEP amplitude in Session 2 correlated with performance following cTBS or sham intervention. The results of these exploratory analyses revealed no significant correlation between baseline cortical excitability and post-stim/sham motor behavior (see supplementary Table S7). Overall, our results suggest that individuals with stronger baseline levels of excitability tended to show greater MEP suppression which was not associated with difference in motor performance.

In conclusion, the current results showed no significant effect of inhibitory cTBS applied to the left M1 prior to schema-mediated motor learning on corticospinal excitability, overall motor sequence execution, retention of learned movement transitions, or schema-mediated integration of novel movement transitions. Our results therefore do not provide evidence for a causal role of the left M1 in schema-mediated motor learning.

## 5. Data availability

The raw data as well as the analyzed data corresponding to the figures presented in the text, and the scripts used to produce them are publicly available on Open Science Framework (https://osf.io/cpqbf).

## 6. Author contributions

**S. Reverberi:** Methodology, Software, Formal analysis, Investigation, Data curation, Writing – original draft, Writing – review & editing, Visualization; **K. Cuypers:** Methodology, Writing – review & editing; **B.R. King:** Conceptualization, Writing – review & editing, Funding acquisition; **G. Albouy:** Conceptualization, Writing – review & editing, Supervision, Project administration, Funding acquisition.

## Acknowledgements / Funding

This study was supported by funding from the FWO Research Foundation Flanders (G0B1419N). GA also received additional support from the FWO Research Foundation (G099516N, G0D7918N, G0B1419N, 1524218N) and internal funds from KU Leuven. SR was supported by a fellowship from the FWO Research Foundation (11C6221N). The authors would like to thank Mareike A. Gann, Melina Hehl, and Sima Chalavi for assistance with data collection.

## 7. Declaration of competing interests

The authors declare that no competing interests exist.

## Notes

https://osf.io/cpqbf

### Competing Interest Statement

The authors have declared no competing interest.

