## Supplementary material for "No evidence for an effect of M1 cTBS on schema-mediated motor sequence learning"

### Random SRTT performance

*Table S1: Results of the statistical analyses of performance on the pseudo-random SRT task measured prior to and following the sequential SRT task in Sessions 1 and 2, respectively, presented in Fig. 2A in the main text. 4 (Block) x 2 (Group) ANOVAs were run per each session and each variable (A: Response Time, averaged across all correct keypresses in each block of practice; B: Accuracy, i.e., percentage of correct keypresses per block of practice). The significant block x group interaction for RT during Session 1 appears to be driven by slower performance on block 1 of the SRTT in the STIM group (see Fig. 2 in main text manuscript); however, pairwise follow-up comparisons revealed that RT was not significantly different across groups on any single practice block. The significant effect of block on accuracy in Session 2 indicates that accuracy increased with random practice across groups (see Fig. 2). Significant values are marked with an asterisk and bold font. Df = degrees of freedom.*

| Effect | df | F | p | Partial $\eta^2$ |
|---|---|---|---|---|
| **A. Response Time** | | | | |
| *Session 1* | | | | |
| Block | 2.03,93.50 | 5.87 | **0.004*** | 0.113 |
| Block x Group | 2.03,93.50 | 4.73 | **0.01*** | 0.093 |
| Group | 1,46 | 0.51 | 0.48 | 0.011 |
| *Session 2* | | | | |
| Block | 3,138 | 0.43 | 0.73 | 0.009 |
| Block x Group | 3,138 | 2.53 | 0.06 | 0.052 |
| Group | 1,46 | 0.01 | 0.94 | 0.000 |
| **B. Accuracy** | | | | |
| *Session 1* | | | | |
| Block | 1.29,59.51 | 1.84 | 0.18 | 0.038 |
| Block x Group | 1.29,59.51 | 1.21 | 0.29 | 0.026 |
| Group | 1,46 | 3.08 | 0.09 | 0.063 |
| *Session 2* | | | | |
| Block | 2.83,130.32 | 3.52 | **0.02*** | 0.071 |
| Block x Group | 2.83,130.32 | 0.83 | 0.48 | 0.018 |
| Group | 1,46 | 2.76 | 0.10 | 0.057 |

### Exploratory analyses on responders vs non-responders

#### Distribution of Session 2 MEP amplitude change in the STIM group

The distribution of the cTBS-induced change in MEP amplitude within the STIM group in Session 2 (i.e., post-pre cTBS intervention) is depicted below in Fig. S1. The distribution is not significantly different from normality (Shapiro-Wilk test statistic=0.97, *p*=0.67). Although such distribution does not suggest a bimodal division of participants in responder and non-responder categories, we performed exploratory analyses to test whether participants who displayed MEP suppression in Session 2 (i.e., a negative MEP change) performed the SRT task differently from those who displayed MEP increase (i.e., a positive MEP change). These groups of participants are referred to as "responders" and "non-responders", respectively, in the analyses that are reported below.

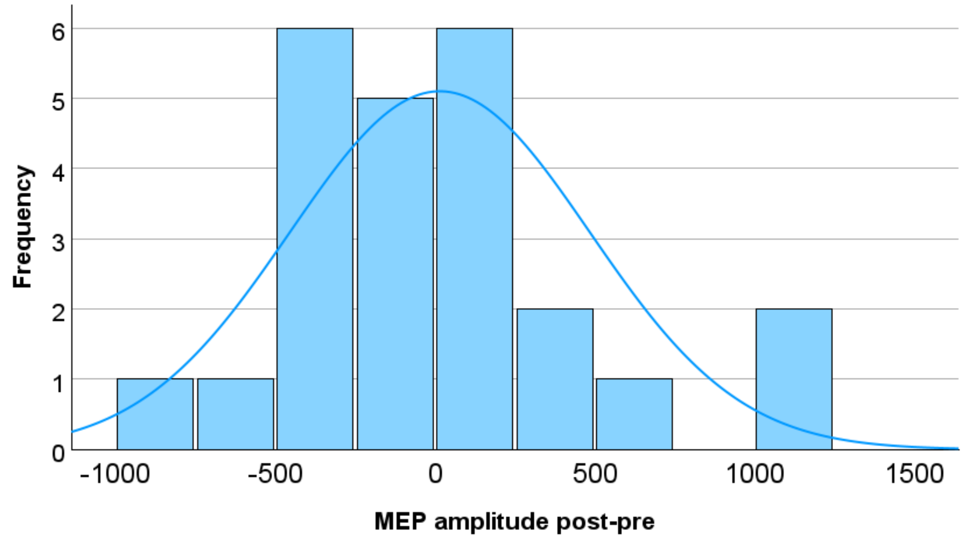


*Figure S1: Distribution of the TBS-induced changes in MEP amplitude in Session 2. Number of bins=9. A normal distribution curve is presented in overlay.*

### Sequential SRTT performance

Sequential SRTT performance in Session 2 for the SHAM group participants (n=24) and for responders (n=13) and non-responders (n=11) from the STIM group are presented in Fig. S2. Performance was compared with repeated-measures ANOVAs using between-subject factor *group* (SHAM/responders/non-responders) and within-subject factor *block*. The results of these analyses are reported below for all transitions (Table S2A), and for novel and learned transitions (Table S2B-C). Results show no significant main effect of group or group x block interaction, suggesting that sequential performance did not differ among groups.

Although no group effects were observed, we also ran follow-up exploratory analyses directly comparing responders and non-responders within the STIM group. No significant group differences in performance were observed, although there was a trend for overall faster performance in responders at test (group effect on response time for all transitions: *p*=0.08, learned and novel transitions: *p*=0.09).

*Table S2: Results of the statistical analyses on the sequential SRTT performance during Session 2. 20/4 (Block) x 3 (Group) ANOVAs were run for each variable (A: Response Time; B: Accuracy). Response Time (RT) and Accuracy measures are provided for all transitions (A) and for learned transitions (B), and novel transitions (C) during Session 2 only. Significant values are marked with an asterisk and bold font. Df = degrees of freedom; Part η2 = partial eta squared; B x G = block x group interaction.*

| | RT | | | | Accuracy | | | |
|---|---|---|---|---|---|---|---|---|
| **Effect** | **df** | **F** | ***p*** | **Part $\eta^2$** | **df** | **F** | ***p*** | **Part $\eta^2$** |
| ***A. All Transitions*** | | | | | | | | |
| *Session 2 Training* | | | | | | | | |
| Block | 4.97,223.54 | 67.78 | **<0.001*** | 0.601 | 10.04,451.86 | 0.88 | 0.55 | 0.019 |
| Group | 2,45 | 0.96 | 0.39 | 0.041 | 2,45 | 0.69 | 0.51 | 0.030 |
| B x G | 9.94,223.54 | 1.46 | 0.16 | 0.061 | 20.08,451.86 | 0.75 | 0.78 | 0.032 |
| *Session 2 Test* | | | | | | | | |
| Block | 3,135 | 0.85 | 0.47 | 0.019 | 3,135 | 2.14 | 0.10 | 0.045 |
| Group | 2,45 | 1.70 | 0.19 | 0.070 | 2,45 | 1.63 | 0.21 | 0.068 |
| B x G | 6,135 | 0.77 | 0.59 | 0.033 | 6,135 | 0.74 | 0.62 | 0.032 |
| ***B. Learned Transitions*** | | | | | | | | |
| *Session 2 Training* | | | | | | | | |

| | | | | | | | | |
|---|---|---|---|---|---|---|---|---|
| Block | 7.20,324.05 | 41.64 | **<0.001*** | 0.481 | 7.94,357.08 | 2.03 | **0.04*** | 0.043 |
| Group | 2,45 | 1.38 | 0.26 | 0.058 | 2,45 | 0.91 | 0.41 | 0.039 |
| B x G | 14.40,324.05 | 1.23 | 0.25 | 0.052 | 15.87,357.08 | 0.83 | 0.65 | 0.035 |
| *Session 2 Test* | | | | | | | | |
| Block | 3,135 | 2.21 | 0.09 | 0.047 | 3,135 | 4.11 | **0.01*** | 0.084 |
| Group | 2,45 | 1.94 | 0.16 | 0.079 | 2,45 | 1.61 | 0.21 | 0.067 |
| B x G | 6,135 | 0.46 | 0.84 | 0.020 | 6,135 | 1.33 | 0.25 | 0.056 |
| ***C. Novel Transitions*** | | | | | | | | |
| *Session 2 Training* | | | | | | | | |
| Block | 4.36,196.19 | 68.42 | **<0.001*** | 0.603 | 11.70,526.28 | 1.24 | 0.25 | 0.027 |
| Group | 2,45 | 0.63 | 0.54 | 0.027 | 2,45 | 0.39 | 0.68 | 0.017 |
| B x G | 8.72,196.19 | 1.55 | 0.14 | 0.065 | 23.39,526.28 | 0.96 | 0.52 | 0.041 |
| *Session 2 Test* | | | | | | | | |
| Block | 2.89,130.10 | 0.66 | 0.58 | 0.014 | 3,135 | 0.30 | 0.82 | 0.007 |
| Group | 2,45 | 1.26 | 0.29 | 0.053 | 2,45 | 1.14 | 0.33 | 0.048 |
| B x G | 5.78,130.10 | 0.94 | 0.47 | 0.040 | 6,135 | 0.44 | 0.85 | 0.019 |

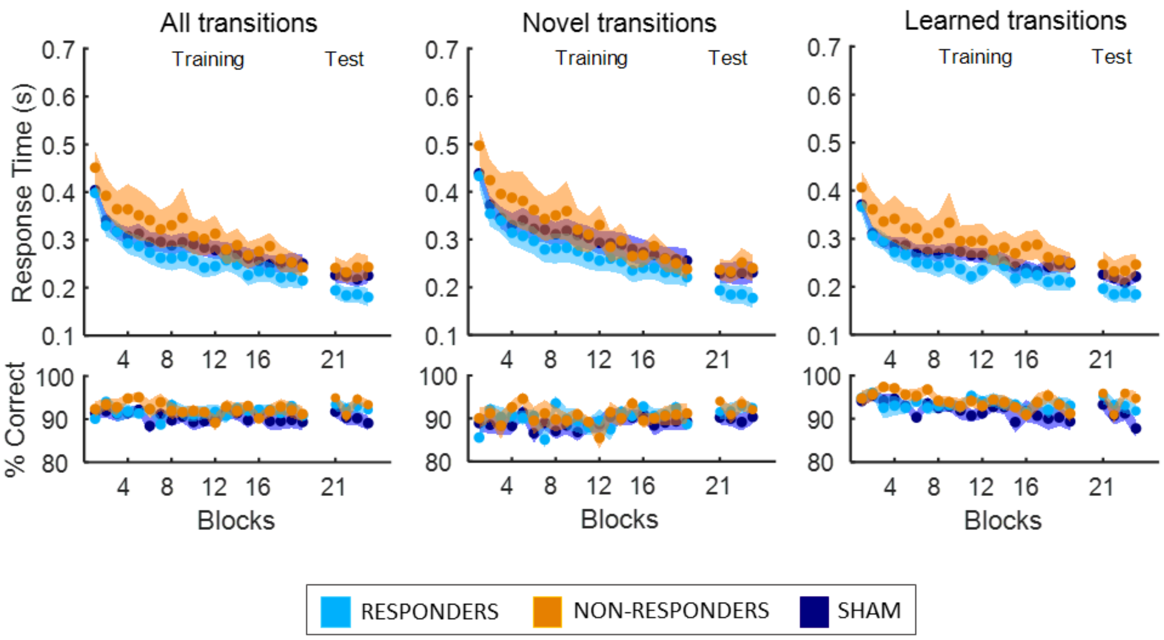


*Figure S2: SRTT performance in Session 2. Mean response time (seconds) and accuracy (% correct transitions) per block are shown separately for the SHAM group (dark blue) and within the STIM group for responders (light blue) and non-responders (orange). Performance averaged across all, novel, and learned transitions. Shaded areas represent the standard error of the mean (SEM).*

### Exploratory analyses on left versus right hand performance

We investigated whether inhibitory cTBS versus sham stimulation of the left M1 affected subsequent bimanual motor sequence performance differently in the left and right hand (ipsilateral and contralateral to the stimulation, respectively). To do so, we contrasted response time for correct keypresses and accuracy, averaged across the keys pressed with the left hand (i.e., keys 1, 2, 3, 4) and with the right hand (i.e., keys 5, 6, 7, 8) during each block of Session 2, between the STIM and SHAM groups (2 *groups* x 20/4 Training/Test *blocks* ANOVAs, as for main text Figure 2). Note that the novel keys (i.e., keys 2 and 3, see experimental procedure and Fig. 1 in the main text) were both pressed by the left hand, so analyses of performance for novel and learned keys were therefore only conducted on left hand data. Results of these analyses are presented in Fig. S3 and Table S3.

In the right hand, there was a significant block x group interaction for the response time measure during training, whereby stimulation resulted in a steeper learning curve on the second half of training as compared to sham stimulation. Note that this effect appeared to be transient as the block by group interaction was no longer observed at post-training test. No group effects were observed on the performance accuracy measure.

In the left hand, no group effects were observed on the response time measure for all keys. However, there was a significant main effect of group on accuracy at test, whereby accuracy in the STIM group was higher than in the SHAM group. We also analyzed performance separately for novel and learned keys performed with the left hand. There were no group effects on response time for either novel or learned keys. There was a significant main effect of group on accuracy at test for novel keys, but not learned keys, indicating that the group effect observed on all left-hand keys was driven by higher accuracy in the STIM group for novel movements. We additionally examined whether learned and novel keys of the left hand were differently performed across the two experimental groups with a repeated-measures ANOVAs with the between-subject factor *group* and the within-subject factor *transition type* (novel vs. learned) on Session 2 test data. The results revealed no interaction of the factors group and transition type, suggesting that learned versus novel keys were not differently performed between groups after extensive task practice (response time – main effect *transition type*: $F(1,46)=0.78$, $p=0.38$, $\eta^2=0.017$; main effect *group*: $F(1,46)=0.21$, $p=0.65$, $\eta^2=0.005$; interaction: $F(1,46)=0.46$, $p=0.50$, $\eta^2=0.010$; accuracy – main effect *transition type*: $F(1,46)=0.00$, $p=0.97$, $\eta^2=0.000$; main effect *group*: $F(1,46)=5.74$, $p=0.02$, $\eta^2=0.111$; interaction: $F(1,46)=0.00$, $p=0.96$, $\eta^2=0.000$). These results therefore suggest that, contrary to our expectations, inhibitory left M1 stimulation - as compared to sham - resulted in an *increase* in performance accuracy on the novel movements performed with the hand ipsilateral to the stimulation.

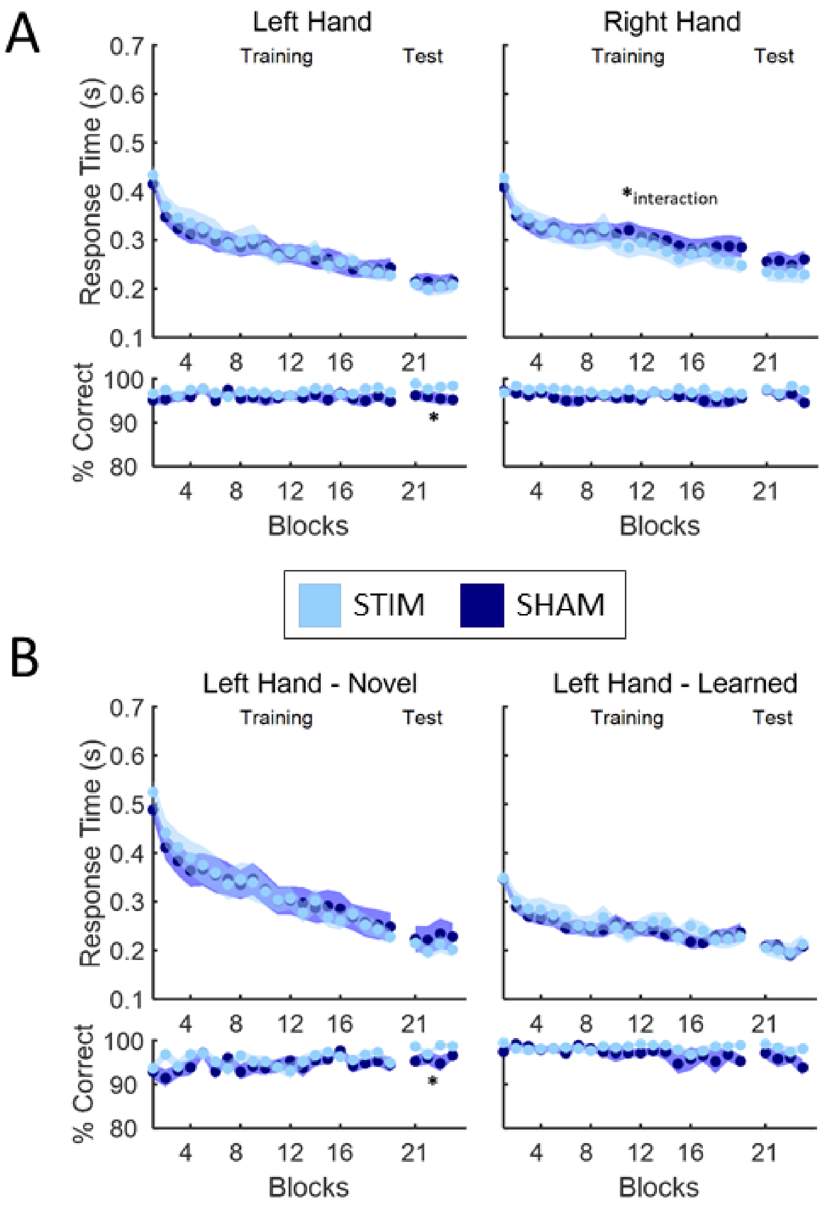


*Figure S3: Performance of the left-hand and right-hand keys in Session 2. Mean response time (seconds) and accuracy (% correct keys) per block are shown separately for the STIM group (light blue) and SHAM group (dark blue). Performance averaged across all movement transitions (A) or separately across learned and novel transitions (B, left hand only). Shaded areas represent the standard error of the mean (SEM). Significant effects are indicated with an asterisk.*

*Table S3: Results of the statistical analyses of sequential SRTT performance in Session 2. Response Time (RT) and Accuracy measures are provided for the for the right hand (B, learned keys only), and for the left hand (B) for all keys (1) as well as learned (2) and novel (3) keys. Significant values are marked with an asterisk and bold font. Df = degrees of freedom; Part η2 = partial eta squared; B x G = block x group interaction.*

| | RT | | | | Accuracy | | | |
|---|---|---|---|---|---|---|---|---|
| Effect | df | F | *p* | Part $\eta^2$ | df | F | *p* | Part $\eta^2$ |
| ***A. Right hand*** | | | | | | | | |
| *Session 2 Training* | | | | | | | | |
| Block | 6.25,287.67 | 43.50 | **<0.001*** | 0.486 | 10.34,475.44 | 1.01 | 0.43 | 0.022 |
| Group | 1,46 | 0.15 | 0.70 | 0.003 | 1,46 | 2.01 | 0.16 | 0.042 |
| B x G | 6.25,287.67 | 2.37 | **0.03*** | 0.049 | 10.34,475.44 | 0.66 | 0.77 | 0.014 |

| | | | | | | | | |
|---|---|---|---|---|---|---|---|---|
| *Session 2 Test* | | | | | | | | |
| Block | 3,138 | 0.29 | 0.83 | 0.006 | 2.72,125.14 | 3.06 | **0.04*** | 0.062 |
| Group | 1,46 | 0.95 | 0.34 | 0.020 | 1,46 | 1.57 | 0.22 | 0.033 |
| B x G | 3,138 | 0.71 | 0.55 | 0.015 | 2.72,125.14 | 1.88 | 0.14 | 0.039 |
| ***B. Left hand*** | | | | | | | | |
| ***1. All keys*** | | | | | | | | |
| *Session 2 Training* | | | | | | | | |
| Block | 5.19,239.10 | 53.25 | **<0.001*** | 0.537 | 9.27,426.30 | 0.71 | 0.70 | 0.015 |
| Group | 1,46 | 0.04 | 0.84 | 0.001 | 1,46 | 1.71 | 0.20 | 0.036 |
| B x G | 5.19,239.10 | 0.97 | 0.44 | 0.021 | 9.27,426.30 | 0.91 | 0.52 | 0.019 |
| *Session 2 Test* | | | | | | | | |
| Block | 3,138 | 0.58 | 0.63 | 0.013 | 3,138 | 0.72 | 0.54 | 0.015 |
| Group | 1,46 | 0.16 | 0.69 | 0.003 | 1,46 | 5.64 | **0.02*** | 0.109 |
| B x G | 3,138 | 0.48 | 0.70 | 0.010 | 3,138 | 0.46 | 0.71 | 0.010 |
| ***2. Learned keys*** | | | | | | | | |
| *Session 2 Training* | | | | | | | | |
| Block | 8.69,399.67 | 19.15 | **<0.001*** | 0.294 | 4.28,196.86 | 1.11 | 0.36 | 0.023 |
| Group | 1,46 | 0.16 | 0.69 | 0.003 | 1,46 | 1.41 | 0.24 | 0.030 |
| B x G | 8.69,399.67 | 1.09 | 0.37 | 0.023 | 4.28,196.86 | 1.11 | 0.36 | 0.024 |
| *Session 2 Test* | | | | | | | | |
| Block | 3,138 | 2.35 | 0.08 | 0.049 | 2.86,131.75 | 1.69 | 0.18 | 0.035 |
| Group | 1,46 | 0.00 | 0.99 | 0.000 | 1,46 | 3.32 | 0.08 | 0.067 |
| B x G | 3,138 | 0.53 | 0.67 | 0.011 | 2.86,131.75 | 0.83 | 0.48 | 0.018 |
| ***3. Novel keys*** | | | | | | | | |
| *Session 2 Training* | | | | | | | | |
| Block | 4.37,201.03 | 57.19 | **<0.001*** | 0.554 | 11.53,530.30 | 1.69 | 0.07 | 0.035 |
| Group | 1,46 | 0.00 | 0.95 | 0.000 | 1,46 | 1.12 | 0.30 | 0.024 |
| B x G | 4.37,201.03 | 1.08 | 0.37 | 0.023 | 11.53,530.30 | 1.18 | 0.30 | 0.025 |
| *Session 2 Test* | | | | | | | | |
| Block | 2.71,124.73 | 1.63 | 0.19 | 0.034 | 2.49,114.53 | 0.62 | 0.58 | 0.013 |
| Group | 1,46 | 0.40 | 0.53 | 0.009 | 1,46 | 5.61 | **0.02*** | 0.109 |
| B x G | 2.71,124.73 | 0.71 | 0.54 | 0.015 | 2.49,114.53 | 1.34 | 0.27 | 0.028 |

### Exploratory analyses on single sequential elements

To explore stimulation effects on specific keys, we compared STIM vs. SHAM group performance on each element in the sequence stream. We averaged response time for correct keypresses and accuracy for all presentations of each sequence element during Session 2 Training and Test runs, then entered the results in a 2 *groups* (STIM/SHAM) x 8 *ordinal positions* (1 through 8, see main text Fig. 1 for details) ANOVAs.

Results are presented in Fig. S4 and Table S4. For both response time and accuracy, there was a significant effect of ordinal position at both Training and Test. We also found a significant group difference in accuracy at Test whereby accuracy was greater in the STIM as compared to the SHAM group. To investigate whether this difference was driven by specific keys, we performed follow-up independent-samples t-tests to contrast STIM vs SHAM accuracy of each sequence element. The results of these follow-up analyses are reported in Table S5. A significant group difference was detected for ordinal position VI, corresponding to novel key 3, with higher accuracy displayed in the STIM compared with SHAM group. However, this effect did not survive Benjamini-Hochberg correction of the False Discovery Rate (applied over the 8 sequence elements). There was no significant effect of group or interaction of group x ordinal position on accuracy at Training, nor on response time at either task run.

Altogether, these results suggest that, while performance differed between fingers (likely due to inherent inter-digit skill differences), this was not affected by the stimulation protocol.

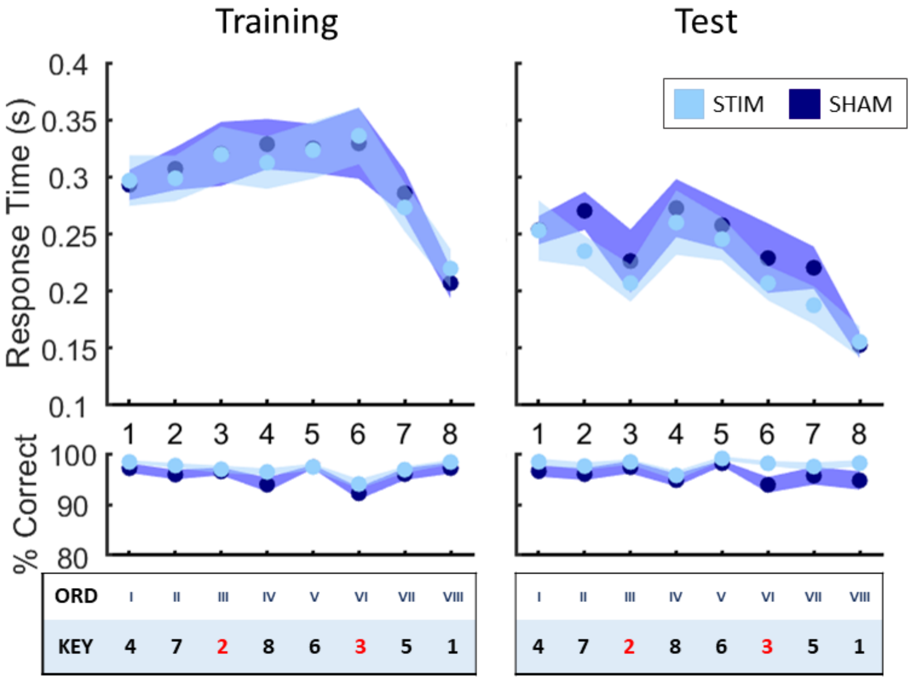


*Figure S4: Performance of each sequential element in Session 2 Training (left) and Test (right) runs. Mean response time (seconds) and accuracy (% correct keys) per ordinal position (1 through 8) are shown separately for the STIM group (light blue) and SHAM group (dark blue). Tables (bottom) indicate the corresponding key pressed at each ordinal position in the sequence (novel keys are presented in red font). Shaded areas represent the standard error of the mean (SEM).*

*Table S4: Results of the statistical analyses of performance on each sequential element in Session 2 (A: Response Time; B: Accuracy). Significant values are marked with an asterisk and bold font. Df = degrees of freedom; Part η2 = partial eta squared; O x G = ordinal position x group interaction.*

| Effect | df | F | p | Partial $\eta^2$ |
|---|---|---|---|---|
| **1. Response Time** | | | | |
| *Session 2 Training* | | | | |
| Ordinal position | 4.02,184.90 | 21.11 | **<0.001*** | 0.315 |
| Group | 1,46 | 0.01 | 0.94 | 0.000 |
| O x G | 4.02,184.90 | 0.35 | 0.85 | 0.007 |
| *Session 2 Test* | | | | |

| | | | | |
|---|---|---|---|---|
| Ordinal position | 4.50,206.92 | 14.28 | **<0.001*** | 0.237 |
| Group | 1,46 | 0.54 | 0.47 | 0.012 |
| O x G | 4.50,206.92 | 0.50 | 0.76 | 0.011 |
| **2. Accuracy** | | | | |
| *Session 2 Training* | | | | |
| Ordinal position | 4.58,210.58 | 12.01 | **<0.001*** | 0.207 |
| Group | 1,46 | 2.03 | 0.16 | 0.042 |
| O x G | 4.58,210.58 | 0.82 | 0.53 | 0.017 |
| *Session 2 Test* | | | | |
| Ordinal position | 5.07,233.35 | 3.03 | **0.01*** | 0.062 |
| Group | 1,46 | 4.18 | **0.047*** | 0.083 |
| O x G | 5.07,233.35 | 0.99 | 0.43 | 0.021 |

*Table S5: Results of independent samples t-tests (STIM vs SHAM) on accuracy during the Session 2 Test run, averaged for each of the 8 ordinal positions (Ord; 1 through 8) in the sequence. Degrees of freedom=46 for all tests. Significant values are marked with an asterisk and bold font. Reported values are uncorrected for multiple comparisons.*

| ORD | t | p | Cohen's d |
|---|---|---|---|
| **I** | 1.45 | 0.16 | 0.42 |
| **II** | 1.15 | 0.26 | 0.33 |
| **III** | 0.76 | 0.45 | 0.22 |
| **IV** | 0.62 | 0.54 | 0.18 |
| **V** | 1.38 | 0.18 | 0.40 |
| **VI** | 2.62 | **0.01*** | 0.76 |
| **VII** | 0.95 | 0.35 | 0.27 |
| **VIII** | 1.79 | 0.08 | 0.52 |

### Exploratory analyses on time elapsed between stimulation and sequential performance

We investigated whether the time elapsed between TMS stimulation (specifically, time of MEP collection prior to cTBS) and the start of sequential SRTT practice in Session 2 was related to motor performance in the STIM group. We detected a significant correlation between time from stimulation and accuracy for novel transitions at training (Table S6); however, this correlation did not survive Benjamini-Hochberg correction of the False Discovery Rate.

*Table S6: Correlations between time from stimulation and: (A) response time in the Session 2 sequential SRTT, (B) accuracy in the Session 2 sequential SRTT within the STIM group. Online gains were calculated as performance difference between block 1 of training and the average of the 4 test blocks. Significant correlations are marked with an asterisk and bold font. Reported values are uncorrected for multiple comparisons.*

| | |
|---|---|
| **A. Response time** | |
| All transitions – training | r=0.06, p=0.79 |
| All transitions – test | r=0.07, p=0.75 |
| All transitions – online gains | r=0.17, p=0.44 |
| Learned transitions – training | r=0.07, p=0.74 |
| Learned transitions - test | r=0.13, p=0.54 |

| | |
|---|---|
| Learned transitions – online gains | r=0.06, p=0.80 |
| Novel transitions – training | r=0.05, p=0.83 |
| Novel transitions – test | r=0.01, p=0.98 |
| Novel transitions – online gains | r=0.22, p=0.31 |
| **B. Accuracy** | |
| All transitions – training | r=-0.39, p=0.06 |
| All transitions – test | r=-0.08, p=0.72 |
| All transitions – online gains | r=-0.30, p=0.16 |
| Learned transitions – training | r=-0.22, p=0.31 |
| Learned transitions - test | r=-0.05, p=0.80 |
| Learned transitions – online gains | r=-0.25, p=0.24 |
| Novel transitions – training | **r=-0.46, p=0.024*** |
| Novel transitions – test | r=-0.08, p=0.71 |
| Novel transitions – online gains | r=-0.24, p=0.27 |

### Exploratory analyses on time of day, age, vigilance, and MEP amplitude

We examined whether the time of day at which the experimental sessions took place, participants' age, and participants' vigilance as assessed with the psychomotor vigilance task (PVT, Dinges & Powell, 1985) and the Stanford Sleepiness Scale (SSS, Hoddes, 1972) were related to baseline MEP amplitude or MEP change during each experimental session.

Participants were tested on average around 1pm [range: 9:26 – 18:07] in both Session 1 (STIM group: 13:52±00:33; SHAM: 13:48±00:29; independent samples t-test t=0.08, *p*=0.94) and Session 2 (STIM: 13:35±00:33; SHAM: 13:16±00:29; t=0.43, *p*=0.67). Over the whole sample, there was no correlation between time of day and baseline MEP amplitude in Session 1 (r=0.09, *p*=0.57) nor in Session 2 (r=-0.17, *p*=0.24) nor between time of day and MEP change in Session 1 (r=-0.17, *p*=0.24) or Session 2 (STIM group only: r=0.15, *p*=0.48).

Participants' age was marginally negatively correlated with baseline MEP amplitude in Session 1 (r=-0.30, *p*=0.04), but not in Session 2 (r=-0.10, *p*=0.51). We also observed no correlation between age and MEP change in either session (Session 1, STIM+SHAM: r=0.19, *p*=0.19; Session 2, STIM: r=0.12, *p*=0.59).

Participants' objective vigilance as assessed with the PVT did not correlate with baseline MEP amplitude in either Session 1 (r=-0.07, *p*=0.64) or Session 2 (r=0.01, *p*=0.97). PVT performance also did not correlate with MEP change over the whole sample in Session 1 (r=-0.08, *p*=0.58) nor within the STIM group in Session 2 (r=0.02, *p*=0.92). Similarly, subjective vigilance as assessed with the SSS did not correlate with either MEP measure in Session 1 (SSS vs baseline MEP: r=-0.15, *p*=0.33; SSS vs MEP change: r=0.06, *p*=0.71) nor in Session 2 (SSS vs baseline MEP, STIM+SHAM: r=0.10, *p*=0.52; SSS vs MEP change, STIM: r=0.35, *p*=0.09).

Altogether, these results suggest that time of day, age, and vigilance were not correlated with baseline MEP amplitude nor MEP amplitude change during either experimental session.

### Exploratory analyses on the relationship between baseline MEP amplitude and learning-induced or stimulation-induced plasticity

To test whether participants who displayed higher baseline cortical excitability were more susceptible to show learning-induced or stimulation-induced plasticity, we examined whether (1) MEP amplitude prior to sequence learning in Session 1 correlated with learning-induced MEP change (over all participants) and (2) MEP amplitude prior to stim/sham intervention in Session 2 correlated with MEP change from post- to pre-intervention (in STIM and SHAM group participants, separately). The results of these analyses are reported in Table S7A. We observed a significant negative correlation over the whole sample between Session 1 MEP amplitude at baseline and MEP amplitude change pre to post motor sequence learning, suggesting that higher baseline MEP amplitude was correlated with greater learning-induced MEP suppression. We also detected a significant negative correlation between cTBS-induced MEP change and baseline MEP amplitude in Session 2 within STIM group participants, suggesting that higher baseline MEP amplitude also correlated with greater cTBS-induced MEP suppression. This correlation was marginally significant in responders (r=-0.56, *p*=0.049) but not in non-responders (r=0.30, *p*=0.37). Although no correlation was detected within SHAM group participants overall, a significant correlation was observed also in SHAM "responders" (i.e., participants who displayed MEP suppression post-sham stimulation; r=-0.75, *p*=0.003) but not in SHAM "non-responders" (r=0.16, *p*=0.65). These results suggest that responders with higher baseline corticospinal excitability were more susceptible to reductions in MEP amplitude irrespective of the stimulation condition (cTBS or sham).

Additionally, we investigated whether baseline MEP amplitude correlated with behavioral performance in either experimental session (Table S7B-C). Overall, we did not observe a correlation between baseline MEP amplitude and behavioral performance in either session. There was a significant negative correlation between baseline MEP amplitude and accuracy on all transitions during the Session 1 test blocks; this was driven by one participant displaying the highest MEP amplitude and lowest accuracy within the sample (removing this participant results in a correlation of r=-0.12, *p*=0.41).

*Table S7: Correlations between baseline MEP amplitude in each experimental session and: (A) MEP changes pre- to post-learning (Session 1) or pre- to post-stim/sham intervention (Session 2), (B) response time in the Session 2 sequential SRTT, and (C) accuracy in the Session 2 sequential SRTT. Online gains were calculated as performance difference between block 1 of training and the average of the 4 test blocks. Significant correlations are marked with an asterisk and bold font. Reported values are uncorrected for multiple comparisons.*

| **A. MEP change** | Session 1 (STIM+SHAM) | Session 2 (STIM) | Session 2 (SHAM) |
|---|---|---|---|
| MEP change | **r=-0.66, p<0.001*** | **r=-0.42, p=0.04*** | r=-0.34, p=0.11 |
| **B. Response time** | | | |
| All transitions – training | r=0.11, p=0.46 | r=-0.12, p=0.57 | r=-0.05, p=0.82 |
| All transitions – test | r=0.04, p=0.81 | r=-0.17, p=0.44 | r=-0.11, p=0.60 |
| All transitions – online gains | r=0.01, p=0.93 | r=-0.06, p=0.77 | r=0.15, p=0.48 |
| Learned transitions – training | | r=-0.18, p=0.40 | r=-0.07, p=0.74 |
| Learned transitions - test | | r=-0.19, p=0.38 | r=-0.12, p=0.57 |
| Learned transitions – online gains | | r=0.03, p=0.90 | r=0.21, p=0.32 |
| Novel transitions – training | | r=-0.07, p=0.76 | r=-0.03, p=0.91 |
| Novel transitions – test | | r=-0.13, p=0.53 | r=-0.08, p=0.70 |
| Novel transitions – online gains | | r=-0.12, p=0.59 | r=0.08, p=0.70 |

| | | | |
|---|---|---|---|
| **3. Accuracy** | | | |
| All transitions – training | r=-0.12, p=0.42 | r=-0.21, p=0.32 | r=0.16, p=0.45 |
| All transitions – test | **r=-0.36, p=0.01*** | r=-0.13, p=0.53 | r=0.10, p=0.66 |
| All transitions – online gains | r=0.08, p=0.61 | r=-0.05, p=0.80 | r=-0.21, p=0.34 |
| Learned transitions – training | | r=-0.21, p=0.32 | r=0.20, p=0.36 |
| Learned transitions - test | | r=-0.23, p=0.28 | r=0.16, p=0.46 |
| Learned transitions – online gains | | r=-0.03, p=0.88 | r=-0.13, p=0.56 |
| Novel transitions – training | | r=-0.17, p=0.42 | r=0.11, p=0.60 |
| Novel transitions – test | | r=-0.02, p=0.95 | r=0.02, p=0.95 |
| Novel transitions – online gains | | r=-0.05, p=0.81 | r=-0.23, p=0.29 |
